# Influenza hemagglutinin drives viral entry via two sequential intramembrane mechanisms

**DOI:** 10.1101/2020.01.30.926816

**Authors:** Anna Pabis, Robert J. Rawle, Peter M. Kasson

**Affiliations:** Department of Cell and Molecular Biology, Uppsala University, Uppsala Sweden; Departments of Molecular Physiology and Biological Physics and of Biomedical Engineering, University of Virginia, Charlottesville VA USA 22908.

## Abstract

Enveloped viruses enter cells via a process of membrane fusion between the viral envelope and a cellular membrane. For influenza virus, mutational data have shown that the membrane-inserted portions of the hemagglutinin protein play a critical role in achieving fusion. In contrast to the relatively well-understood ectodomain, a predictive mechanistic understanding of the intramembrane mechanisms by which influenza hemagglutinin drives fusion has been elusive. We have used molecular dynamics simulations of fusion between a full-length hemagglutinin proteoliposome and a lipid bilayer to analyze these mechanisms. In our simulations, hemagglutinin first acts within the membrane to increase lipid tail protrusion and promote stalk formation and then acts to engage the distal leaflets of each membrane and promote stalk widening, curvature, and eventual fusion. These two sequential mechanisms, one occurring prior to stalk formation and one after, are consistent with experimental measurements we report of single-virus fusion kinetics to liposomes of different sizes. The resulting model also helps explain and integrate prior mutational and biophysical data, particularly the mutational sensitivity of the fusion peptide N-terminus and the length sensitivity of the transmembrane domain. We hypothesize that entry by other enveloped viruses may also utilize sequential processes of acyl tail exposure followed by membrane curvature and distal leaflet engagement.

## Introduction

Enveloped viruses infect cells via a process of membrane fusion that releases the viral core into the cytoplasm. For influenza virus, the viral hemagglutinin is the sole protein component necessary and sufficient for fusion (1–3). Mature hemagglutinin consists of two subunits, one of which mediates receptor binding and the other fusion (4, 5). The fusion subunit HA2 consists of a transmembrane anchor, a soluble ectodomain that refolds to bring host and target membranes together (6), and an amphipathic fusion peptide that inserts into cellular membranes to help mediate fusion (7). Mutational data show that viral fusion peptides do not simply anchor the ectodomain in the cellular membrane but must act within the host membrane for the virus to successfully achieve fusion (8–11). The nature of this intramembrane activity has been hotly debated, and emerging evidence suggests that it likely differs from endogenous vesicle fusion machinery in the cell (12). These mechanisms are difficult to resolve experimentally because fusion is mediated by a loose, transient complex of viral proteins that have been activated for fusion. Here, we use molecular dynamics simulations of fusion by full-length influenza hemagglutinin assemblies to show two sequential intramembrane activities of influenza fusion peptides required for fusion. This model integrates and explains prior experimental data on potential fusion peptide mechanisms and yields a broad integrative view of the process from membrane apposition to fusion pore opening.

Viral membrane fusion is believed to proceed through a series of lipidic intermediates (13). Current understanding, including recent structural results (14–16), is summarized as follows. Initial membrane apposition brings host and target membranes together with patchy hydration at the interface but no gross changes in bilayer structure (17). The first structural intermediate involving membrane change is the formation of a fusion stalk where the proximal apposed membrane leaflets begin to merge, forming a small region of continuous aliphatic density across the intervening polar layer (18, 19). This stalk then expands into a hemifusion diaphragm where proximal leaflets are merged but distal leaflets remain intact. Fusion pore opening represents the final topological change in fusion, resulting in continuous water density between the lumens of the virus and the target. There is relatively strong consensus about these intermediates. Substantial uncertainty remains, however, regarding the precise role of influenza fusion proteins in promoting these intermediates, their fine structure, and any off-pathway states in fusion.

Many orthogonal approaches have yielded data regarding potential roles for influenza fusion peptides in mediating membrane fusion, but no single mechanism has been able to explain all the available data to date. We highlight a few findings here that are of particular relevance to this work. Different mutations to the N-terminal fusion peptide of hemagglutinin (HA2 subunit after proteolytic cleavage) can either result in complete arrest of fusion or can permit lipid mixing but impair content transfer through a fusion pore (8, 9, 11). Such impairment can occur even in the presence of ectodomain refolding to the post-fusion conformation (8, 9). These phenotypes have been reproduced in both cells expressing full-length hemagglutinin and infectious virus, and although phenotypes vary between the two systems (20), the data strongly support an intramembrane role for fusion peptides in driving fusion. Simulations have found that these fusion peptides can induce acyl tail protrusion in proportion to their ability to induce lipid mixing (21) and that this acyl tail protrusion is key to formation of stalk-like intermediates on the fusion pathway (22–24). It has been suggested spectroscopically that this such activity may correlate with semi-closed fusion peptide conformations (25). Experimental studies on PEG-mediated vesicle fusion with exogenous fusion peptides offer indirect support for this model (26). Separately, spectroscopic experiments and simulations on isolated influenza fusion peptides suggest that they are capable of inducing membrane curvature (27–30), which could alter the free-energy barriers to fusion either by changing the starting reference state or by stabilizing highly curved intermediates (31, 32). However, as we show in this manuscript, the overall starting curvature of the target membrane does not detectably affect the rate of influenza virus lipid mixing.

Testing these theories directly has been challenging because biochemical manipulations feasible on isolated peptides are challenging on full-length hemagglutinin and even more so on infectious virus. Additionally, because the membrane context of these transient fusion assemblies is critical, fusion phenotype can vary even between cells expressing hemagglutinin and *bona fide* virions (20). We have used molecular dynamics simulations to approximate the fusion of a pH-activated influenza virion to a planar bilayer, the most common experimental geometry used to measure viral fusion kinetics at high spatiotemporal resolution (33–36). We employ a multi-resolution approach: we use atomic-resolution simulations to capture fusion stalk formation and pore opening (Fig. 1) and coarse-grained simulations for the slower evolution of hemifusion intermediates between these events (Fig. 2). We employ perturbation tests to validate our simulations and then compare them against existing or new experimental data not used to tune the simulations.

**Figure 1.**
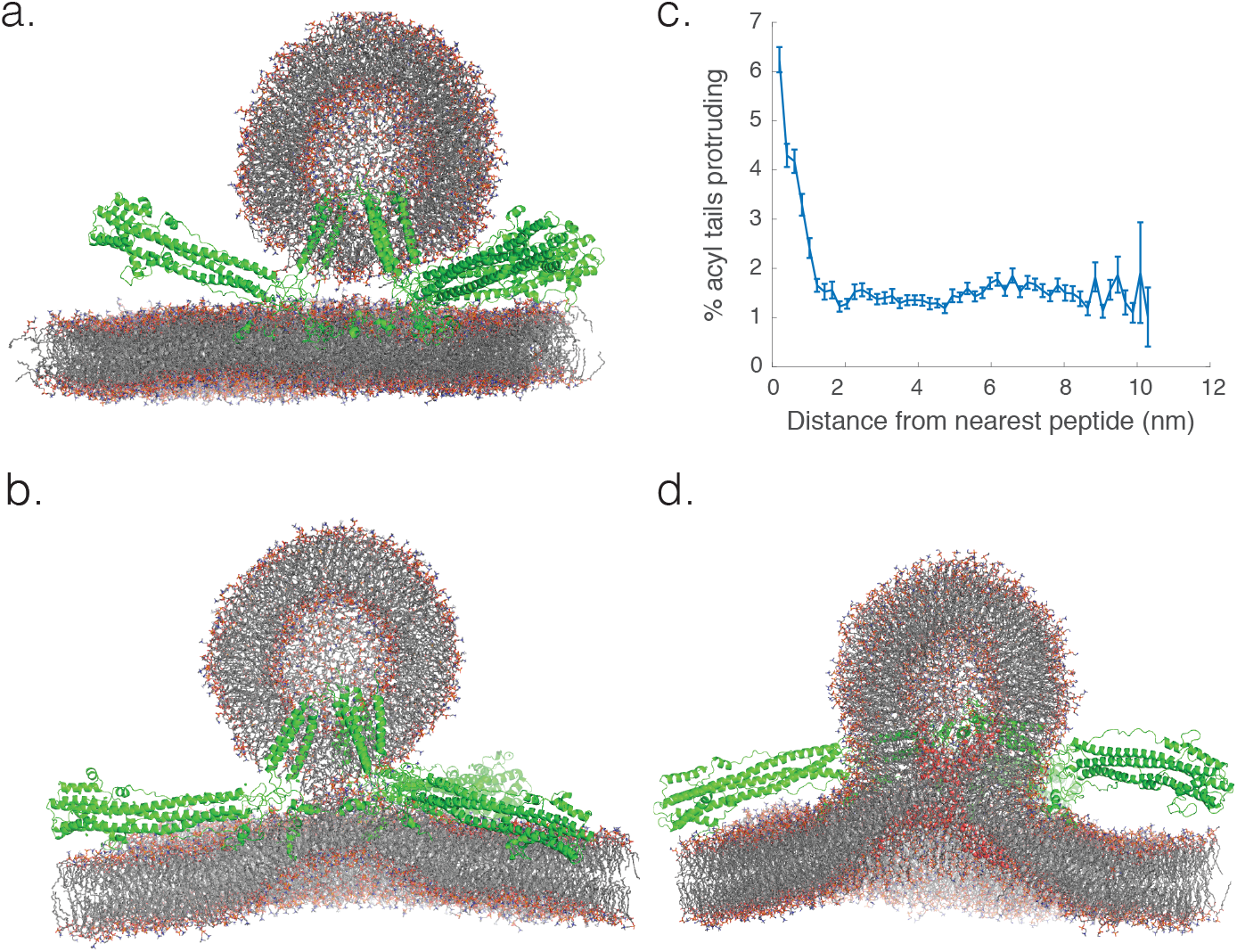
Influenza fusion peptides promote acyl tail protrusion leading to stalk formation. Atomic-resolution simulations of a small proteoliposome docked to a planar bilayer by three full-length hemagglutinin trimers (a) resulted in fusion stalk formation (b) with a rate of 1.7 μs^−1^, estimated over 50 independent simulations starting with the hemagglutinin ectodomains in the postfusion conformation. Fusion peptides promoted local protrusion of phospholipid acyl tails (c), contributing to stalk formation as previously described. A nascent fusion pore is rendered in (d), with water lining the pore in sphere form. Renderings show cutaway of phospholipids in stick form and hemagglutinin proteins in cartoon form, with explicit water and ions not rendered. All three hemagglutinin trimers are present in all simulations. Error bars represent 90% confidence intervals calculated via bootstrap resampling. Water density before and after pore formation is rendered in Fig. S3.

**Figure 2.**
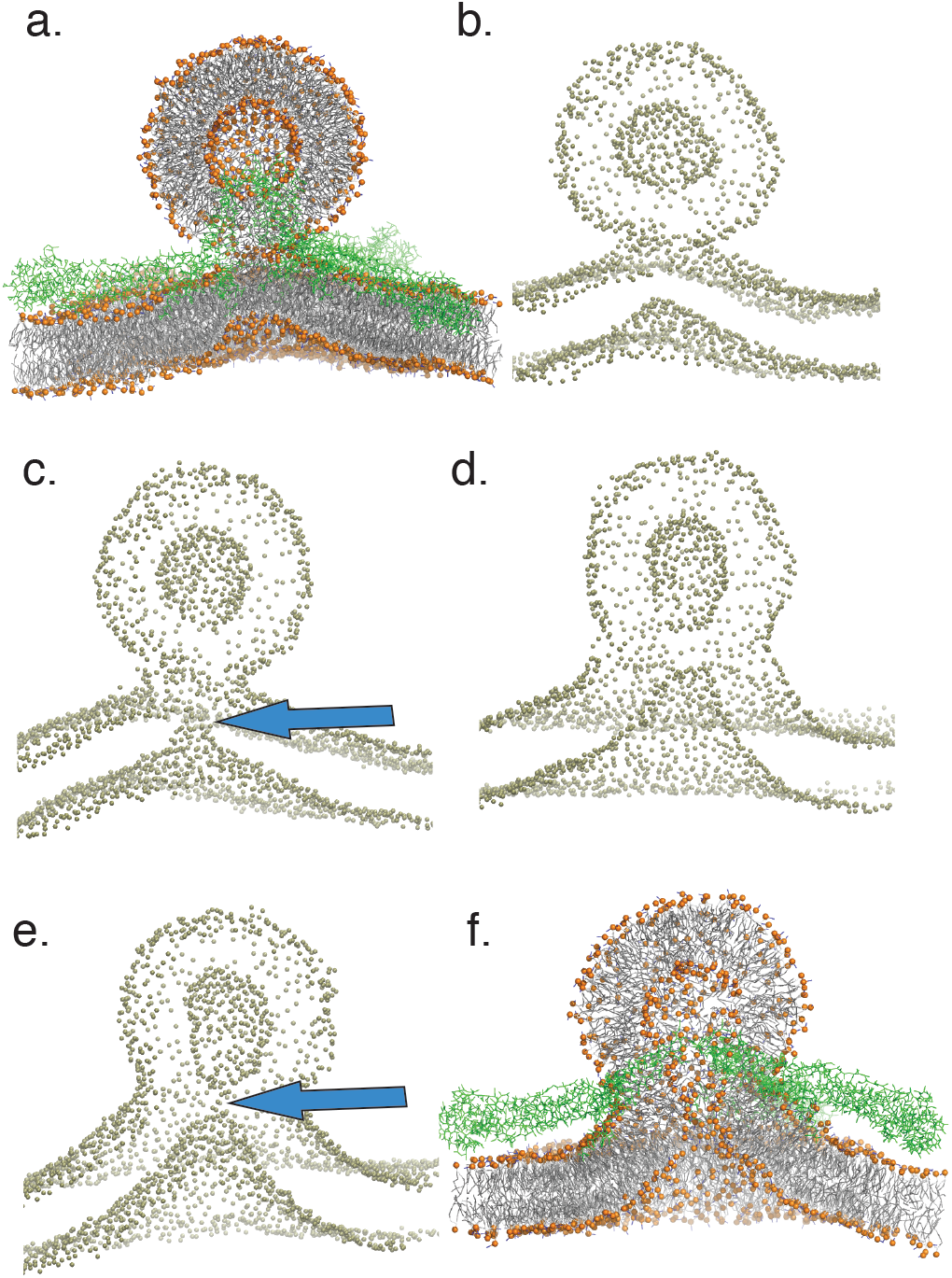
Distal leaflet involvement in the junctional complex leads to membrane bending and fusion pore formation. Starting from a stalk structure (a), cutaway renderings of phospholipid phosphate groups show changes in membrane geometry from the initial stalk (b) to distal leaflet movement towards the junctional complex, here shown at 80 ns of coarse-grained simulation (c). This permits subsequent stalk widening and hemifusion diaphragm formation between the two distal leaflets, rendered at 500 ns of coarse-grained simulation (d) and finally fusion pore formation, rendered at 6.78 μs of coarse-grained simulation (e), and shown again in coarse-grained detail (f). Membrane bending is a key feature of progression to fusion pore formation but primarily occurs after initial stalk formation. Coarse-grained simulation times to pore formation ranged from 267 ns to 6.8 μs after stalk formation. As shown in the following figures, the sequence of events rendered here was a common feature of all simulations leading to fusion pore opening.

## Results and Discussion

All simulations contained a proteoliposome approximating the virus, three full-length hemagglutinin trimers, and a planar lipid bilayer approximating the target membrane. Fusion stalk formation was assessed by analyzing 50 independent atomic-resolution simulations started from a docked state after ectodomain refolding. The timing of early lipidic intermediates in fusion versus ectodomain refolding has not been fully resolved, although it is believed that refolding precedes fusion (16, 37, 38). We therefore chose a starting state consistent with this hypothesis where the energetic contribution from ectodomain refolding has been completed, but where fusion would arrest if not for intramembrane contributions from hemagglutinin. The latter conclusion is demonstrated by fusion peptide mutants that lead to ectodomain refolding but no lipid mixing (11). Simulations were then coarse-grained at the point of stalk formation and run at multiple peptide protonation states, lipid compositions, and bilayer sizes to analyze the determinants of fusion pore formation. To test the robustness of hemagglutinin intramembrane fusion mechanisms to membrane curvature, simulations were also performed using a proteoliposome of twice the diameter (30 nm) and a correspondingly larger bilayer. Finally, simulations were converted back to atomic resolution shortly before fusion pore opening to capture atomic details of the fusion pore. This multi-resolution strategy was chosen because single-water-layer effects in stalk and pore formation particularly benefit from atomic resolution simulations (17).

Simulated fusion stalk formation by full-length hemagglutinin proceeded largely as expected from prior simulations of lipid-only vesicle fusion and of isolated fusion peptides (21–24). Close membrane apposition was rapidly formed, accompanied by patchy dehydration of the interface. Fusion stalks were nucleated by stochastic encounter of protruding acyl tails, promoted by the fusion peptides (Fig. 1). This is qualitatively similar to the emerging understanding of SNARE-mediated fusion stalk formation (39). Using 50 individual simulations, we estimate the rate for stalk formation in this system from an activated, docked conformation at 1.7 μs^−1^ using a Poisson approximation (See Methods). This is in accordance with the expectation that fusion peptide release and hemagglutinin ectodomain refolding are rate-limiting for the process of lipid-mixing. We observed fusion peptides to be conformationally plastic (Fig. S1), visiting conformations similar to each of prior NMR-derived conformational structures (40–42). These conformations were metastable but underwent exchange on the sub-microsecond timescale, while ectodomain structure was well preserved (Fig. S2).

Membrane bending was observed to be a critical feature of fusion, but, unexpectedly, this effect was manifest only in stages subsequent to stalk formation. Simulations showed minor bending of the target bilayer well prior to stalk formation, but this was not rate-limiting for stalk formation. Indeed, the upward displacement of the bilayer was less than one bilayer thickness and not significantly greater when stalks were formed than in time-matched non-productive encounters (p>0.8, Mann-Whitney U test). In contrast, substantial bending occurred in all cases prior to fusion pore formation (Fig. 2), after stalk expansion and leading to direct apposition of the distal leaflets in a hemifusion diaphragm. Thus, for hemagglutinin-mediated fusion of a small proteoliposome with an approximately planar bilayer, we predict that membrane bending affects the rate of fusion pore opening but not the rate of initial stalk formation.

This finding is consistent with single-virus experiments on influenza fusion with small unilamellar vesicles. Virus was bound to GD1a-containing vesicles tethered to a passivated surface inside a microfluidic flow cell as previously described (36), fusion was triggered by a buffer exchange to pH 5, and lipid-mixing kinetics were assessed by the waiting time between pH drop and fluorescence dequenching of fluorescently labeled lipids in the viral envelope. When this procedure was repeated for liposomes extruded at different sizes, lipid-mixing kinetics were not detectably different (Fig. 3). This suggests that lipid-mixing kinetics are not demonstrably altered by changes to target-membrane curvature and deformability. Although alternate explanations are possible, these results are consistent with our simulation predictions that membrane bending primarily occurs after stalk formation, the molecular event that permits stable lipid mixing.

**Figure 3.**
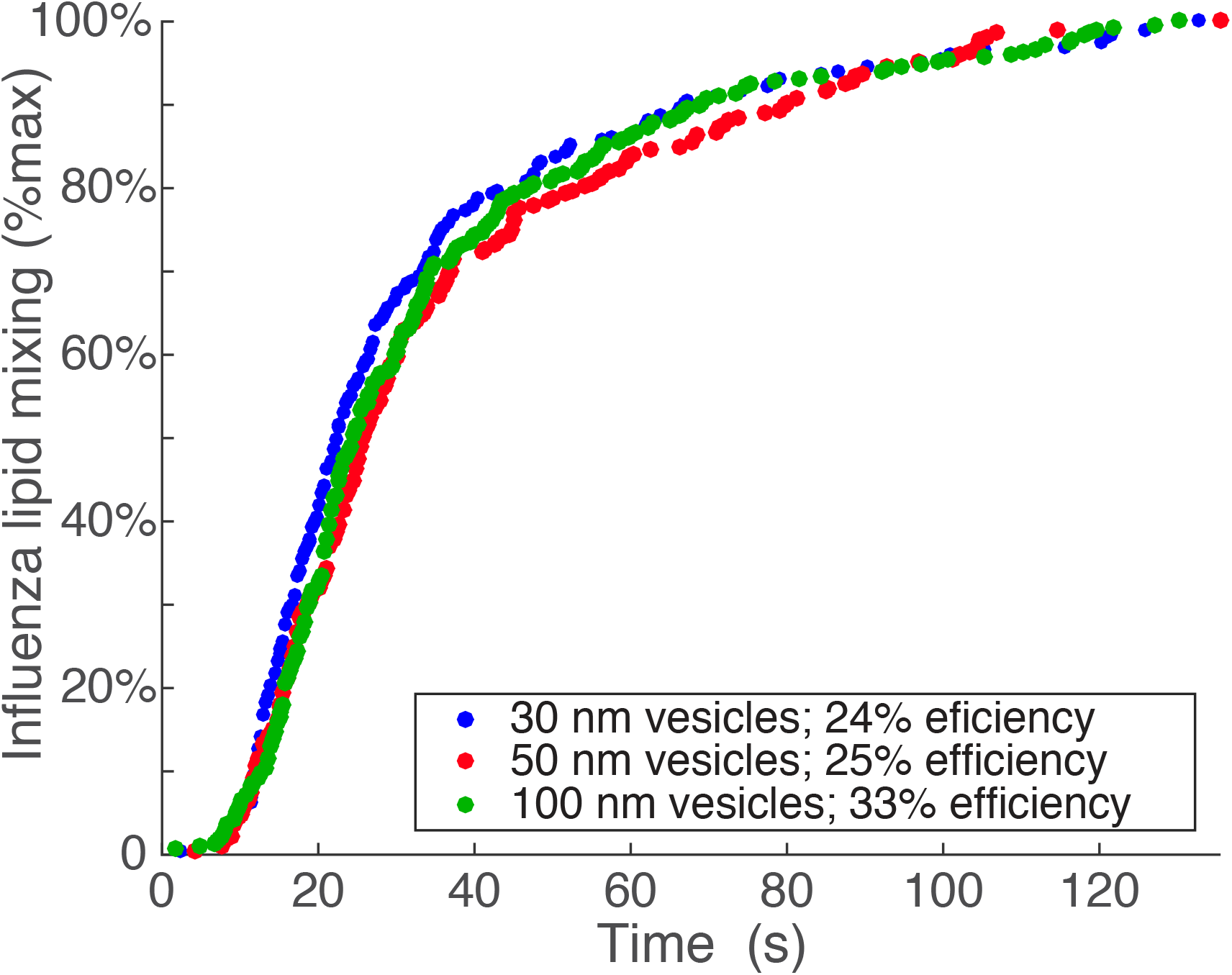
Rates of lipid mixing are not detectably different between influenza virus and target liposomes of different sizes. Single-virus fusion experiments between fluorescently labeled X-31 (H3N2) influenza virus and liposomes extruded at different sizes show no detectable dependence of lipid-mixing rate on vesicle size. This is consistent with the hypothesis that membrane bending primarily occurs after stalk formation. Cumulative distribution curves for individual fusion event distributions are plotted versus time after pH drop. Vesicle sizes refer to the diameter of the extrusion pore used.

In simulations, expansion of the stalk junctional complex and formation of a hemifusion diaphragm (Fig. 2) required three components. These were: an increase in stalk radius, inward movement of the distal leaflets into the junctional complex (Fig. 2c), and subsequent increased membrane bending. Stalk radius expansion occurred simultaneous to inward movement of the distal leaflets (Fig. 4). When the lower leaflet of the bilayer was restrained from upward movement into the stalk complex, no stalk expansion and no fusion pore opening occurred (Fig. S4), suggesting that this upward movement is required for progression to fusion. This multi-step progression to fusion involves several intermediate states; nonetheless a rough estimate of overall timescale can be obtained by treating the opening of a fusion pore from the stalk state as a Poisson process. Under that approximation, the rate of fusion pore formation for small proteoliposomes is 0.18 μs^−1^, estimated from 20 simulations. The general principle of distal leaflet involvement is similar to prior simulations of SNARE-mediated fusion (43), although the details of the membrane complexes and mechanisms by which involvement is achieved differ here.

**Figure 4.**
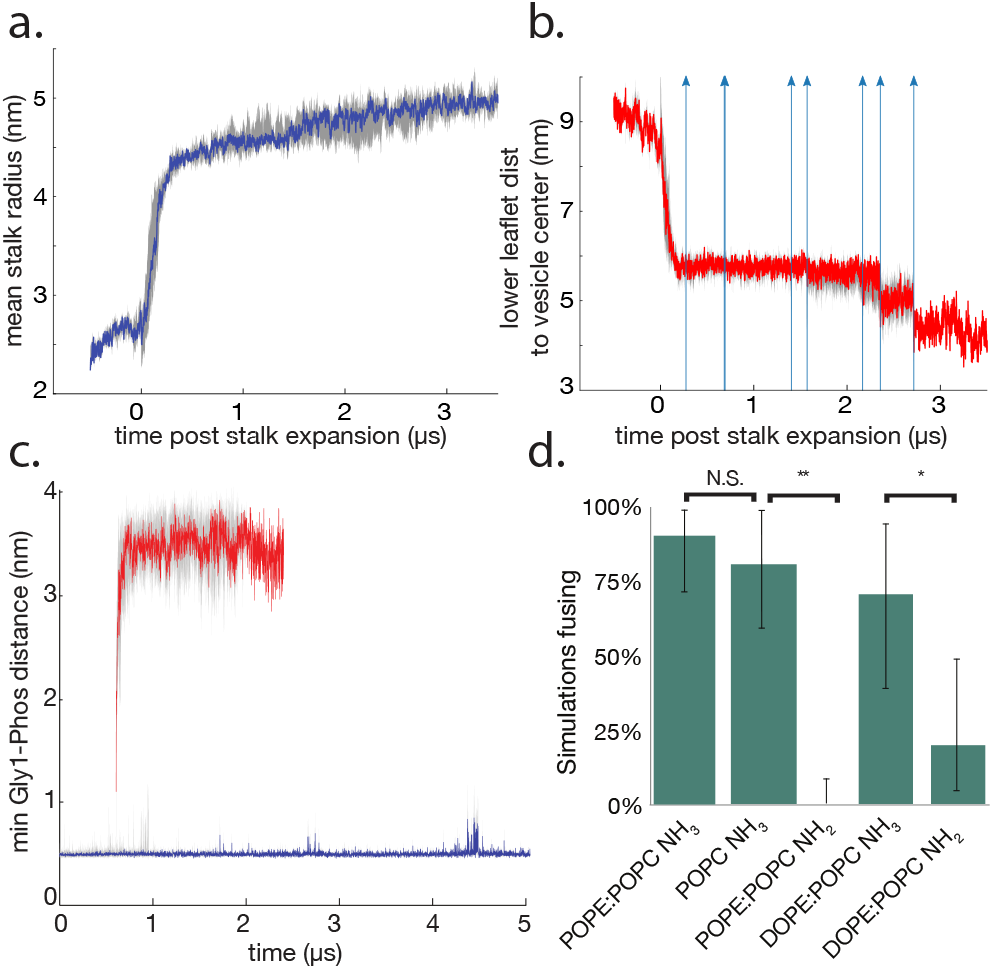
Stalk widening occurs simultaneous to distal-leaflet inward movement and permits later stages of fusion. Stalk radius is estimated via root mean squared distance from the geometric center of the stalk and plotted versus time in panel (a). Stalk expansion occurs simultaneous to inward movement of the bilayer distal leaflet, plotted as minimum distance between the lower leaflet and the vesicle center in panel (b). Quantities are plotted as median value for all unfused trajectories with interquartile range shown in gray. Trajectories are synchronized to the time of stalk expansion (denoted time 0). Fusion pore opening events are denoted by vertical arrows in panel (b). Gly1 to lower-leaflet phosphate distances are plotted in panel (c) and are maintained stably at 0.49 nm (90% CI 0.46-0.55 nm) and when lost rapidly decay to approximately the bilayer thickness (median 3.47 nm, 90% CI 2.91-3.87 nm). In all fusing trajectories in the original dataset, loss of Gly1-Phos contact occurs only after stalk expansion, and commitment to fusion occurs relatively early (Fig. S9). Traces show median and interquartile range of Gly1-Phos distance since simulation start (blue) or loss of contact (red). Panel (d) shows the results of several perturbations: changing the lipid composition to 100% POPC while maintaining protonated peptide N-termini, maintaining the POPE:POPC lipid composition while neutralizing the peptide N-termini. Bars show 90% confidence intervals calculated via bootstrap, and changing the lipid composition to 75% POPC, 25% DOPE with either protonated or neutral N-termini. N.S. denotes not significantly different; ** denotes p < 1e-4, and * denotes p < 0.02 via bootstrapped hypothesis testing. Analogous plots to panels (a-c) for neutral N-termini are given in Figure S10.

Facilitating such inward movement of the distal leaflets appears to be a key role of hemagglutinin. Hemagglutinin fusion peptides localize to the rim of the expanding stalk and may assist the curvature changes associated with stalk widening, but such localization alone was not enough to drive efficient fusion in our simulations in the absence of distal leaflet involvement. In our simulations, occasional deep insertion of a fusion peptide N-terminus resulted in contact with the distal leaflet phosphate. Deeper insertion has been previously suggested to correlate with fusion activity of influenza (44) and HIV (45) fusion peptides. The Gly1-PO4 contact we observed was highly efficient in drawing the distal leaflet into the junctional complex and facilitating progression to fusion. This activity depended on the protonation state of the Gly1 amino terminus: when the terminal NH2 was held uncharged, fusion was greatly reduced (Fig. 4d). Such activity was particularly important in simulations of a smaller periodic patch of bilayer, which causes an increase in the free energy required for upward deformation by some Δz compared to a larger periodic patch (Fig. S5). When we repeated fusion simulations using the same stalk structure and a larger target bilayer, stalk widening and ultimate fusion pore opening were observed both with and without Gly1-PO4 contact, although to a greater extent with a protonated amino terminus (Fig. 5). The same effect was observed in simulations of hemagglutinin-mediated fusion of a vesicle of twice the radius and a correspondingly larger bilayer: 14/20 simulations with a protonated peptide N-terminus proceeded from a stalk state through membrane curvature and stalk widening, while only 1/20 simulations with neutral N-termini showed such progression (Fig. S6). Together, this suggests that trans-bilayer contact by hemagglutinin, when it occurs, can reduce the activation free energy for fusion stalk widening by drawing the distal leaflets into the junctional complex.

**Figure 5.**
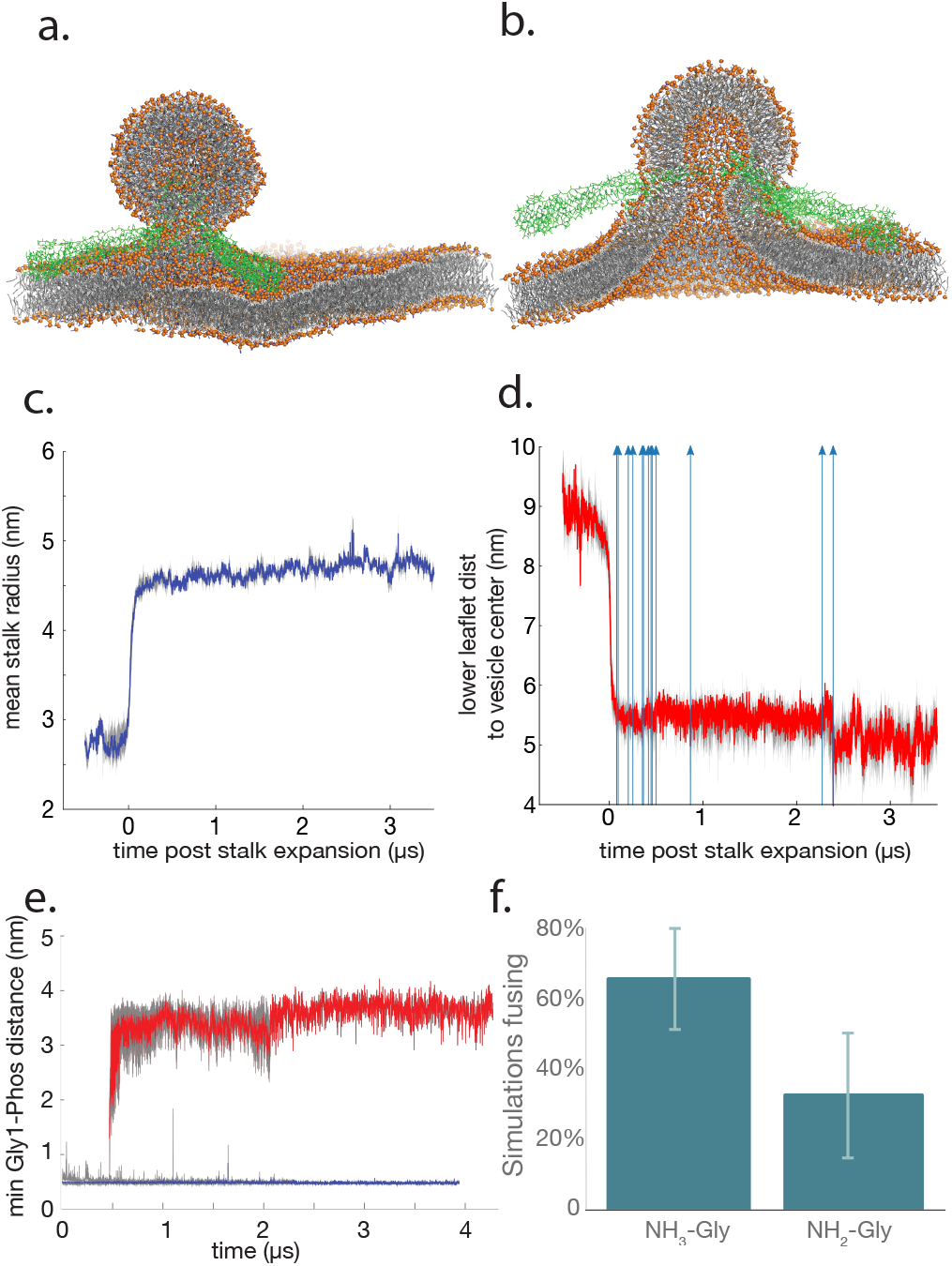
Fusion mechanism is preserved but requirement for Gly1-PO_4_ contact relaxed in simulations of a larger bilayer patch. Fusion was simulated of the same stalk structure as in Figure 4 except with a larger target bilayer. The initial stalk state and a representative fusion pore are rendered in panels (a) and (b) respectively. Plotted in panel (c) is stalk radius expansion estimated as the root mean squared distance of stalk lipids from the center, synchronized to time of expansion. Panel (d) shows the inward movement of the bilayer lower leaflet into the functional complex, and panel (e) shows Gly1-lower-leaflet-phosphate contact distances similar to Figure 4. Data plotted in panels d-e are the aggregate of simulations with protonated and unprotonated glycines. The fraction of these simulations fusing is plotted in panel (f); unlike with a smaller bilayer patch, a substantial fraction of simulations fuse without protonated glycines and without Gly1-Phos contacts. As in Figure 4, data are plotted as median values with interquartile ranges represented in gray; bars in panel (d) show bootstrapped 90% confidence intervals, and blue arrows in panel (b) show times of fusion. The bootstrapped p-value for difference between the mean fusion outcomes is 0.04. Data are plotted for NH2-Gly and NH3-Gly simulations separately in Fig. S11.

These findings may also help explain the requirement for a ≥17-residue hemagglutinin transmembrane anchor but relative mutational tolerance observed experimentally (46–49). In our simulations of a highly curved liposome, the transmembrane domain may facilitate recruitment of the liposome inner leaflet into the stalk complex (Fig. S7), in some cases apparently pulling the liposome inner leaflet into the junctional complex. Such an effect would be further accentuated in a 75-nm diameter viral particle and can indeed be seen in simulations of a 30-nm proteoliposome (Fig. 6) from a stalk state through fusion. In the less-curved membrane of a full-size influenza virus, the activity of the transmembrane anchor may indeed resemble that which we observe for deeply inserted fusion peptides. In neither case, however, does curvature and fusion occur due to force transduction through the fusion peptide or transmembrane domain after stalk formation: when the peptide bonds connecting either the fusion peptide or the transmembrane anchor to the ectodomain were severed after stalk formation, simulations still proceeded to fusion.

**Figure 6.**
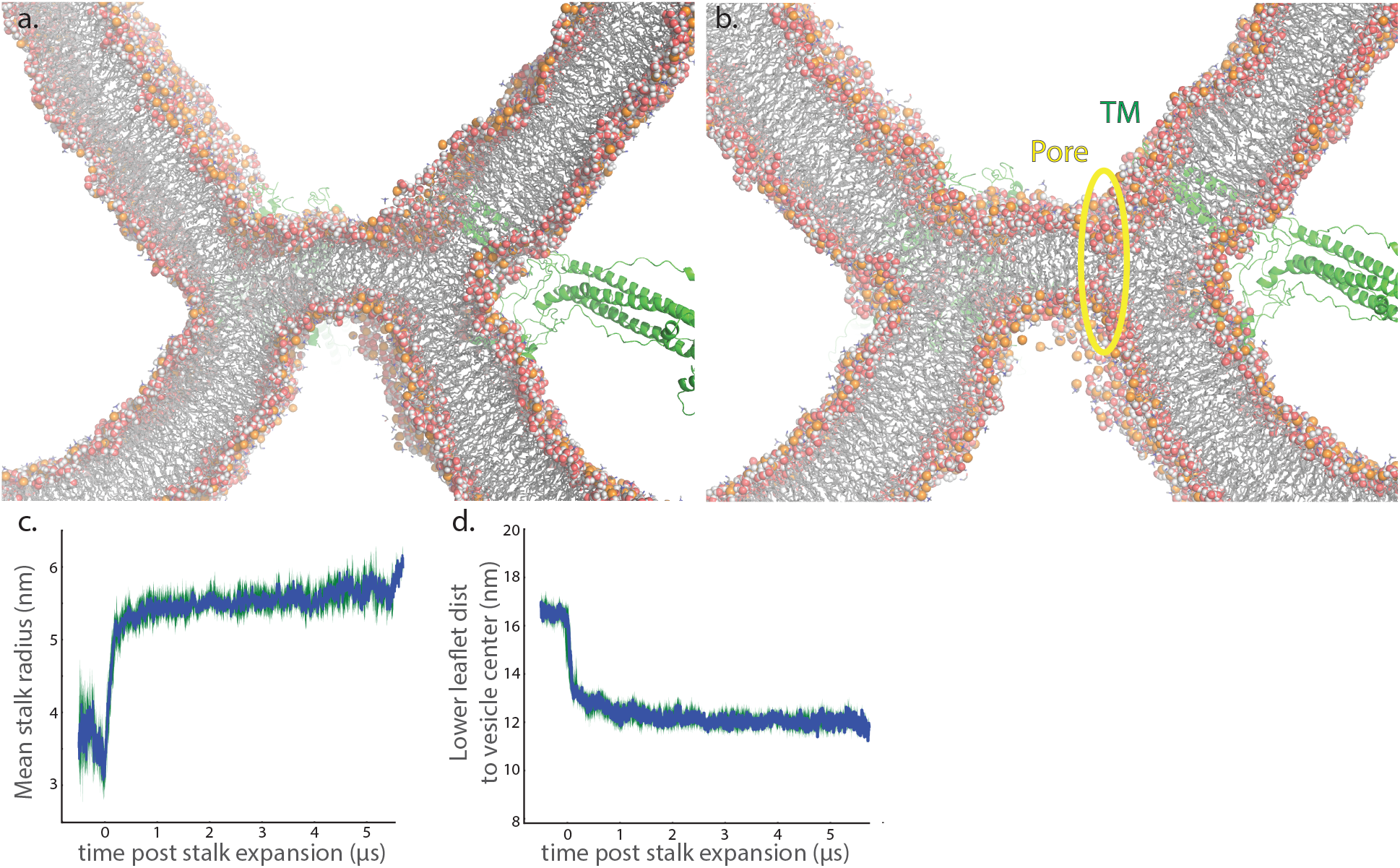
Fusion pore formation. Rendered are snapshots 1ns before (a) and at the time of (b) fusion pore formation between a 30-nm proteoliposome and a 1900 nm^2^ bilayer, simulated at atomic resolution. Full system snapshots of these fusion simulations are shown in Figs. S15 and S16. Stalk radius expansion in preceding coarse-grained simulations of the 30-nm proteoliposome:bilayer system is plotted in panel (c), and the recruitment of the lower leaflet into the hemifusion complex is plotted in panel (d). The fusion pore is nucleated in a region of thinned bilayer and consists of continuous water density spanning the hemifusion diaphragm. Other local water defects that do not achieve continuous transbilayer water density do not nucleate fusion pores. The hemagglutinin transmembrane domains likely contribute to fusion pore formation by anchoring the ends of the hemifusion diaphragm and keeping it thinned.

We also performed a set of simulations to examine the process of fusion pore formation in atomic detail. A coarse-grained simulation snapshot 10 ns prior to fusion pore opening was converted to atomic resolution and 30 simulations started from that point. 67% of these simulations fused, with a median time to fusion of 21 ns, indicating that these late hemifusion structures were highly committed to fusion pore opening in both coarse-grained and atomistic representations. In addition, 10 atomic-resolution simulations were started from three different coarse-grained hemifusion snapshots using a proteoliposome of twice the size (diameter 30 nm). 50% of these larger simulations fused within 50 ns; the lower fusion rate can be attributed to a single coarse-grained snapshot that had likely not yet progressed to be fusion-capable. The process of pore formation was marked by a thinning of the hemifusion diaphragm, followed by formation of a partial water defect in this diaphragm. This partial defect was then followed by water penetration from the opposite side to form continuous water density across the hemifusion diaphragm (Figs. 6, S8). This was rapidly followed by reorientation of the adjacent headgroups to form a pore lined by polar moieties. In our simulations, pore formation occurred in a region of hemifusion diaphragm thinning; no portion of the hemagglutinin protein directly contacted the nascent pore. The initial pore formed in simulated hemagglutinin-driven fusion was thus entirely lipidic in nature. Simulations of pore formation between a 30-nm proteoliposome and a target bilayer suggest that the hemagglutinin transmembrane domain may play a role in pinning the edges of the thinned hemifusion diaphragm (Fig. 6), thus lowering the free energy barrier to pore formation. Mutational evidence suggests a further role for hemagglutinin in fusion pore widening (49, 50); whether this proceeds via protein action in the pore itself or outside the immediate pore region remains unclear.

## Conclusions

Here we report mixed-resolution simulations of influenza membrane fusion between either 15-nm or 30-nm proteoliposomes approximating the virus and a planar bilayer. This system was chosen to mimic several aspects of single-virus fusion experiments within a computationally tractable framework. It simulates the process of membrane fusion after membrane apposition achieved by refolding of the hemagglutinin ectodomains. Experimental evidence is mixed on whether full refolding of all three ectodomains is required for fusion (3, 38, 51–53), but 1) it should constitute the full contribution of the ectodomains towards fusion and 2) fusion peptide mutations have shown that in absence of sufficient intramembrane hemagglutinin activity the refolding of these ectodomains alone is insufficient for fusion (8–11). One key aspect of virus-membrane fusion experiments that our simulations do not include is the presence of cholesterol. We expect that cholesterol will modulate membrane properties but that the fundamental mechanisms elucidated here will remain. Our simulations are thus devised to probe the intramembrane activity of hemagglutinin, particularly the fusion peptide. Because the hemagglutinin proteoliposome simulated is highly curved, the simulations may not fully capture the contribution of hemagglutinin transmembrane domains to fusion. We are, however, able to hypothesize a mechanism of action for the transmembrane domain by close observation of the domain placement during our simulations and by analogy from fusion peptide activity.

Our simulations thus predict two sequential mechanisms for the intramembrane activity of influenza hemagglutinin leading to fusion: first increasing lipid acyl tail exposure and catalyzing stalk formation and second helping recruit distal leaflets into the stalk junctional complex leading to hemifusion diaphragm formation and fusion pore opening. Fusion peptides may also contribute to proximal leaflet curvature (27) and stalk radius expansion: our simulations show that the peptides are well placed for such an effect but do not directly test whether they contribute to fusion in this way. Deep insertion of the fusion-peptide N-terminus, as predicted by these simulations, may be a rare event that contributes directly to membrane fusion, or it may reflect a physical mechanism that is in fact utilized by the transmembrane domain and helps to explain the length requirement for the transmembrane domain observed experimentally. Thus, our findings yield hypotheses for how both intramembrane portions of influenza hemagglutinin contribute to fusion and viral entry. They help explain how mutations to the N-terminus of the fusion peptide and deletions in the transmembrane anchor impair fusion pore opening because those portions could be responsible for distal leaflet recruitment. The multi-resolution simulation approach employed here further yields a platform to examine other contributions to fusion, such as more complex lipid compositions, the potential for fusion peptide-transmembrane domain complexes in the hemifused state, and the effect of different fusion peptide mutations on stages of fusion beyond stalk formation. Such simulations can both help explain the disparate experimental findings on influenza membrane fusion and suggest additional experiments to test the mechanistic hypotheses. More broadly, this paradigm of acyl tail exposure facilitating stalk formation and then distal leaflet recruitment as a key permissive factor for stalk expansion and fusion pore opening may also describe entry by other enveloped viruses. Class I viral fusion proteins would be expected to have the greatest mechanistic similarity to influenza, but testing the generality of these conclusions to entry mediated by class II and class III will help illuminate the mechanistic conservation versus diversity in viral entry.

## Methods

### Simulations from docked state

A hemagglutinin proteoliposome was generated by assembling a model of full-length H3 hemagglutinin corresponding to the X-31 influenza strain. The full-length hemagglutinin model was assembled using Modeller (54) and supplying a crystallographic structure of the postfusion HA2 ectodomain (55), an NMR structure of residues 1-20 of the HA fusion peptide in lipid micelles (40), and modeling the transmembrane domain as an ideal alpha helix. The fusion peptides were initially inserted in the bilayer membrane approximating the membrane depths determined by EPR (40), and the proteoliposome was placed at 16 Å closest approach distance to the bilayer with the hemagglutinin ectodomains positioned between. The positions of the membrane-inserted fusion peptides and transmembrane helices were held fixed, and Modeller was used to generate the structure of the linker and the orientation of the ectodomain via template-based modeling using the templates specified above. The resulting assembly was then relaxed via energy minimization and molecular dynamics simulation (see Supplementary Methods and Fig. S12). The proteoliposome and bilayer were separated by a water layer that was of approximately bulk density (Fig. S13) and displayed liquid-like properties with slowed diffusion (Fig. S13), characteristic of fluid water near a membrane interface but not yet displaying the glassy properties of water between two tightly apposed bilayers (56). The liposome was taken from previous vesicle fusion simulations and had 15-nm diameter (17), and the fusion peptides were inserted into a 2000-lipid bilayer. Different NMR studies have yielded multiple conformational models for the hemagglutinin fusion peptide (41, 42, 57, 58), and our simulation trajectories sampled structures close to all of the pH 5 structures (Fig. S1). Lipid compositions used were POPE:POPC 75:25 and 100% POPC, as indicated. Simulations were run using Gromacs (59) and AMBER99SB-ILDN (60, 61) and lipid parameters we have previously published (17). Detailed parameters are given in the Supplementary Methods. 50 independent simulations were run for 200-500 ns each; simulations and lengths are tabulated in Table S1.

### Simulations from stalk state

Atomic-resolution stalk structures were converted to coarse-grained, and coarse-grained protein parameters were generated from the hemagglutinin conformations in the stalk structures using the *backwards* (62) and *martinize* utilities respectively. Simulations were re-solvated, equilibrated, and restarted using the MARTINI parameters (63) in Gromacs (see Supplementary Methods and Fig. S14 for details). 10 simulations each were run from this stalk state with protonated glycine N-termini and neutral N-termini, 5 with protonated N-termini and POPC lipids, 10 each with protonated and neutral N-termini using 75:25 POPC:DOPE lipids in the target bilayer, and 2 additional control simulations with protonated N-termini and NPT conditions. Simulations and lengths are given in Table S1.

### Simulations of fusion pore opening

A structure was selected 10 (coarse-grained) ns prior to first fusion pore opening in MARTINI simulations and converted to atomic resolution in the CHARMM36 force-field (64, 65) using the *backwards* utility. This was used to start 30 independent simulations (19 with protonated fusion peptide N-termini and 11 with neutral fusion peptide N-termini; trajectories tabulated in Table S1). These were again solvated in TIP3P water with 150 mM NaCl. At start, no fusion pore was present. Simulations were run in Gromacs with parameters identical to the CHARMM simulations from the docked state above.

### Simulations of larger bilayer patches

An additional 30 coarse-grained simulations were performed from the stalk state with a larger bilayer patch (1000 nm^2^), generated as described in the Supplement. 15 of these used protonated N-termini and 15 used neutral N-termini. Run lengths are included in Table S1.

### Simulations of larger vesicles and bilayers

A further 40 coarse-grained simulations were performed from the stalk state using a vesicle of twice the diameter (30 nm) and a corresponding bilayer patch of 1930 nm^2^, generated from stalk states formed by 15-nm vesicles as described in the Supplement. 20 simulations used protonated N-termini and 20 used neutral N-termini. Three of the simulations using protonated N-termini were converted to atomic resolution shortly before pore opening and a total of 10 runs from those structures were performed as described above and in the Supplement. The atomic-resolution system size was 8 million atoms. Simulation run lengths are included in Table S1.

### Single-virus fusion experiments

Labeling of virus, preparation of vesicles and microfluidic flow cells, microscopy, and analysis were performed as previously described (36) with the exception of the extrusion pore size. Briefly, X-31 influenza virus (A/Aichi/68, H3N2) was labeled with Texas Red-DHPE at a self-quenching concentration and allowed to bind to small unilamellar vesicles produced via extrusion of a 65.5% POPC, 20% DOPE, 10% cholesterol, 2% GD1a, and 0.5 Oregon Green-DHPE mixture through pores of the indicated size. The vesicles were immobilized by DNA-lipid binding to a passivated glass coverslip inside a microfluidic flow cell prior to virus binding. After virus binding, unbound virus was washed away by exchange of >20 volumes of buffer. Fusion was initiated by buffer exchange to pH 5.0. Fusion events were detected by dequenching of Texas Red dye in single-virus spots, and the single-event waiting times were compiled into cumulative distribution curves for each fusion condition. Analysis code is available at https://github.com/kassonlab/micrograph-spot-analysis.

### Simulation analysis methods

Distances were measured using Gromacs; data processing was performed in Python. Stalk-formation rates were estimated via a previously published Poisson event model (66, 67); details are given in the Supplement. Stalk expansion was estimated as the root-mean-squared distance of all stalk lipid particles from the geometric center of the stalk, median-filtered over 3-ns windows. Lipid tail protrusion was measured via a custom utility written for Gromacs using definitions specified previously (24). Stalk formation and fusion pore formation were analyzed using alpha complexes in a variation on previous work (68). Details are given in the Supplement, and code for both is available from https://github.com/kassonlab/fusion-shape-analysis.

## Acknowledgements

This work was funded by R01 GM098304 and a Wallenberg Academy Fellowship to P.M.K. Computing time was provided by Folding@Home, the PDC Center for High Performance Computing supported by the Swedish National Infrastructure for Computing, and NSF allocations on Blue Waters and Frontera. Marta Domanska assisted with initial setup of some simulations. We thank many colleagues for helpful discussions, including J. Hays, S. Haldar, A. Villamil Giraldo, C. Stroupe, L. Tamm, L. Chernomordik, and J. Zimmerberg.

