## Supplementary figures and tables for "Influenza hemagglutinin drives viral entry via two sequential intramembrane mechanisms"

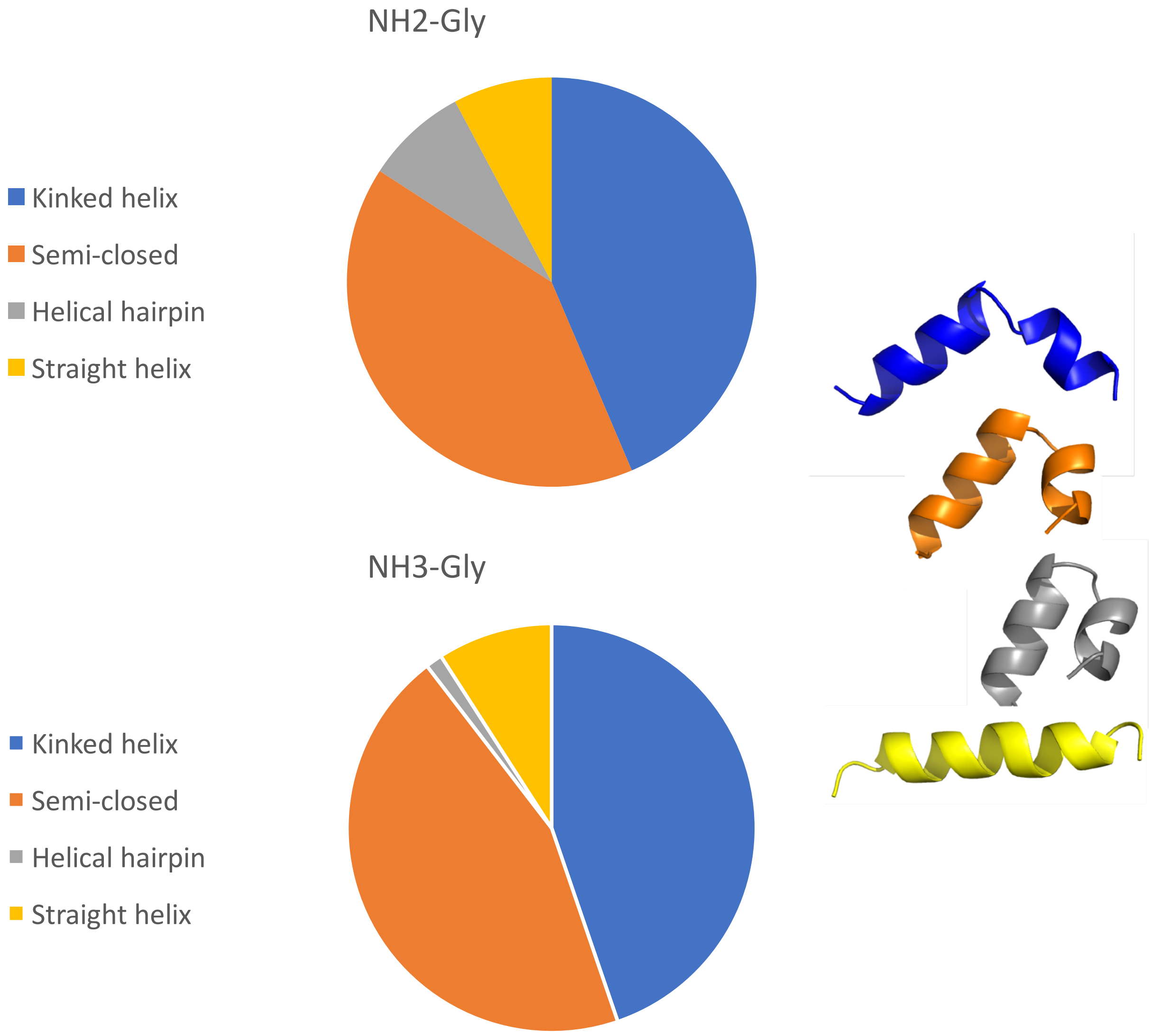


**Figure S1. Conformational states of simulated fusion peptides.** Fusion peptide coordinates were extracted at 200-ps intervals in atomistic simulations, and the aligned RMSD of backbone atoms was measured to each of four conformational models: a kinked helix (1), a semi-closed structure (2), a helical hairpin (3), and a straight helix. The best-fit structure was assigned to each snapshot and the resulting distribution of states calculated for simulations both with a protonated N-terminus ($NH_3$-Gly) and a neutral N-terminus ($NH_2$-Gly). Similar to prior work (4), the sampling here did not detect a substantial difference in the conformational state between protonated and neutral N-terminal states, with the possible exception of the helical hairpin conformation. More extensive sampling of the joint equilibrium between protonation state and insertion depth would be required to detect this more sensitively. Renderings show residues 1-20 in the simulations with minimum RMSD to each reference model, with N-termini on the left.

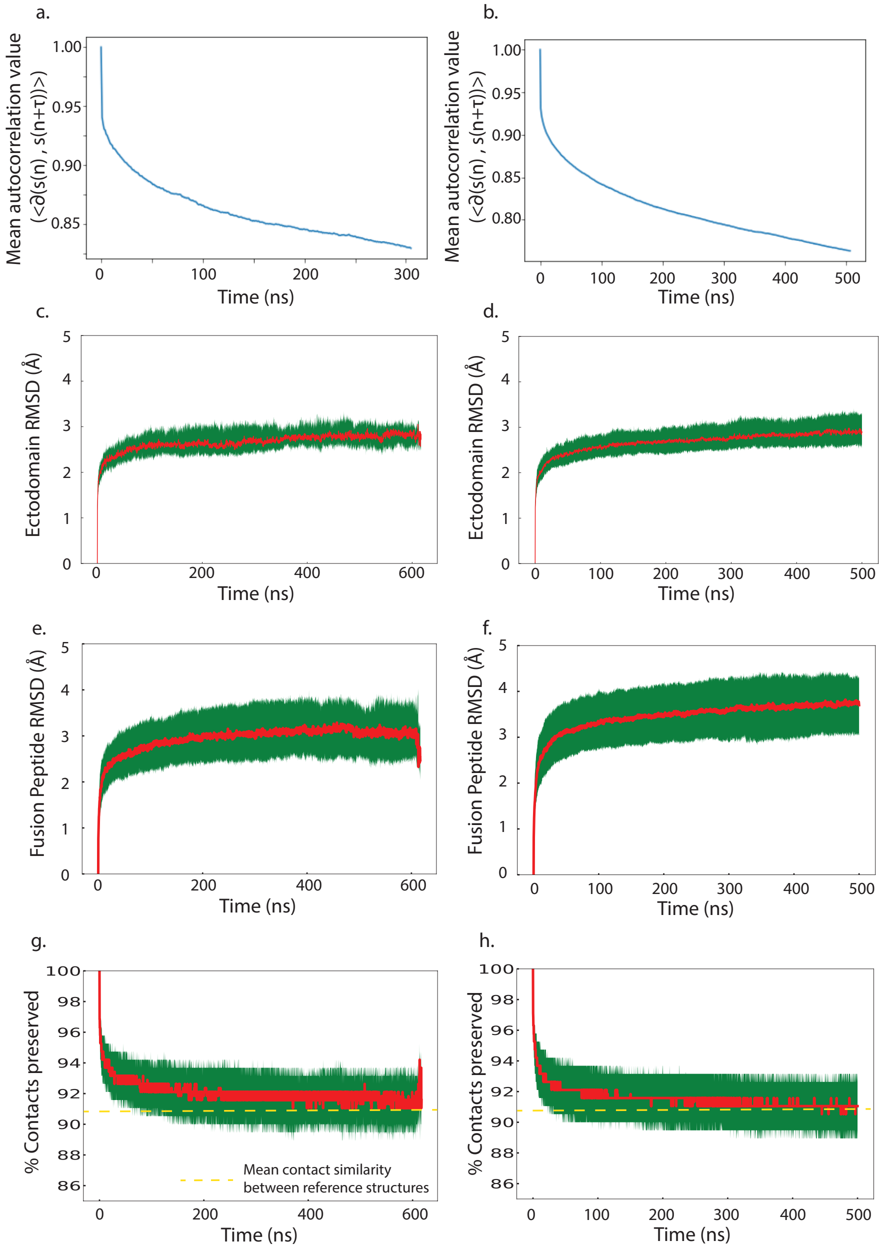


**Figure S2. Time-series analysis of fusion peptide conformations and ectodomain stability**. Time-autocorrelation values are plotted for the conformational states of the fusion peptide in panels (a) and (b) for atomistic simulations with protonated and unprotonated N-termini, respectively. Time-autocorrelation functions are defined as $<\partial(s(n), s(n+\tau)>$, where s(n) is the conformational state of the peptide at time n, using conformational states as defined as in Figure S1. $\partial$ denotes the Kronecker delta function, and $<>$ is averaging across all fusion peptides simulated. Further analysis of conformation-state time series using a Hidden Markov Model with 4 states and lifetimes of 82, 79, 66, and 34 ns respectively, all shorter than the length of the simulations (medians 238 ns for protonated and 500ns for neutral N-terminus simulations, Table S1). In both Markov State Model and Hidden Markov Model analysis (5, 6), all transitions between conformational states were well sampled. Panels (c) and (d) plot alpha-carbon root mean squared deviation (RMSD) over the course of the simulations for protonated and unprotonated N-termini respectively. Panels (d) and (e) plot RMSD for the fusion peptide, and panels (e) and (f) plot fraction of preserved contacts for residues 1-20 of the fusion peptide. Horizontal lines show the mean difference between reference structures. The ectodomain, defined here as the residues 35-178, residues that were well-ordered in two thirds of the post-fusion crystal structure by Wiley and colleagues (7), remained stable over the course of the simulations. Values plotted are median and interquartile range. To assess longer-term stability, protonated simulations were extended to median 477 ns, and these were used to calculate RMSD values.

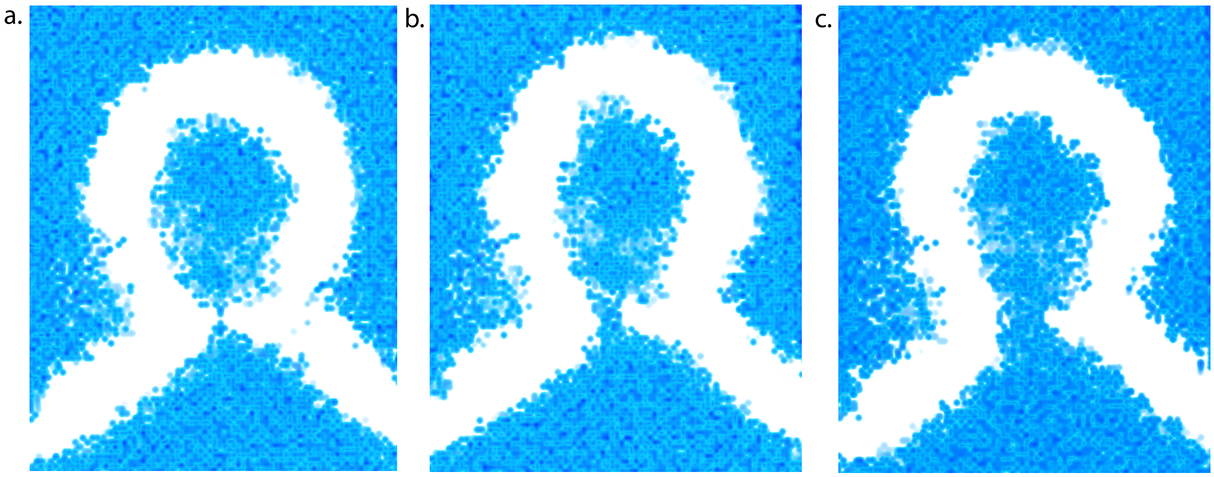


**Figure S3**. **Water density before and after fusion pore formation**. Water density was calculated in voxels 2Å on a side and rendered for a 20-nm slice through the 15-nm proteoliposome. Renderings are shown 2 ns prior to pore formation, where the water density is not yet continuous (**a**), at time of fusion pore formation (**b**), and 4 ns after pore formation (**c**).

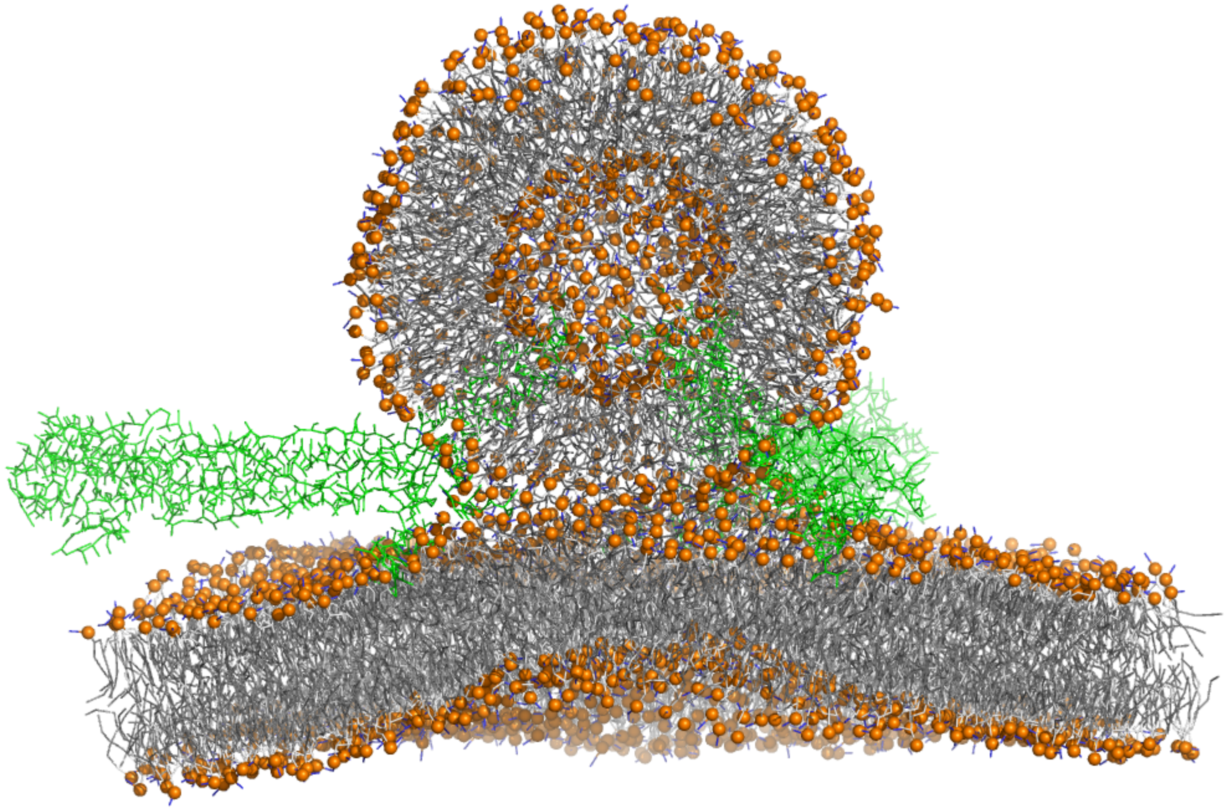

**Figure S4. Involvement of the distal bilayer leaflet is required for stalk expansion and fusion**. Coarse-grained simulations were run from a stalk starting state where lipids in the distal leaflet of the target bilayer were restrained from bending upwards. Lipids were selected as those in the lower leaflet and within 2.5 nm of a fusion peptide, and a spring constant of 250 kJ $mol^{-1}$ $nm^{-2}$. With this minimal set of 116 lipid atoms restrained, stalk expansion was greatly impaired, although minor asymmetric stalk widening was observed on the 1.5 μs timescale, rendered here.

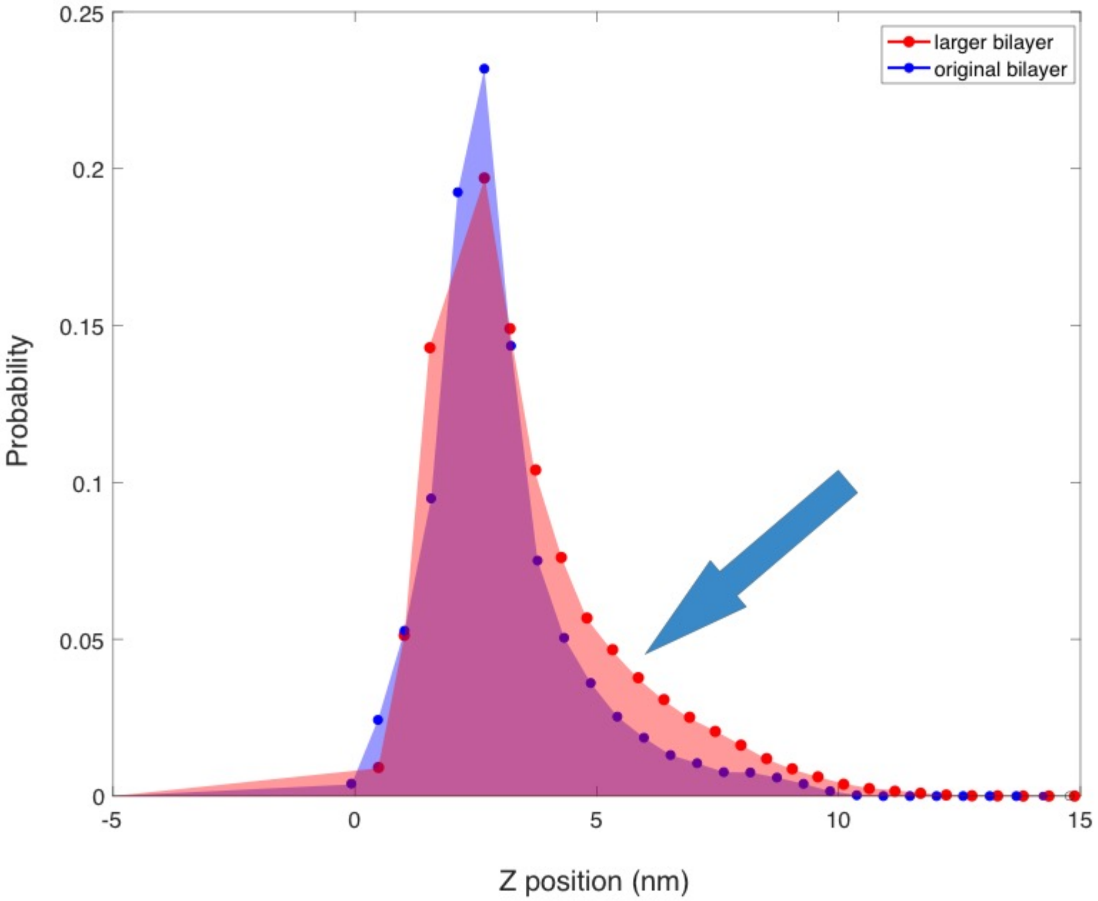


**Figure S5. Target membrane deformation and periodic cell size**. Plotted are histograms of lower-leaflet phosphate position over the first 500ns of a simulation with the original bilayer size (blue) and the larger bilayer size (red). As expected, a larger bilayer size results in greater magnitude fluctuations in z (as indicated by blue arrow).

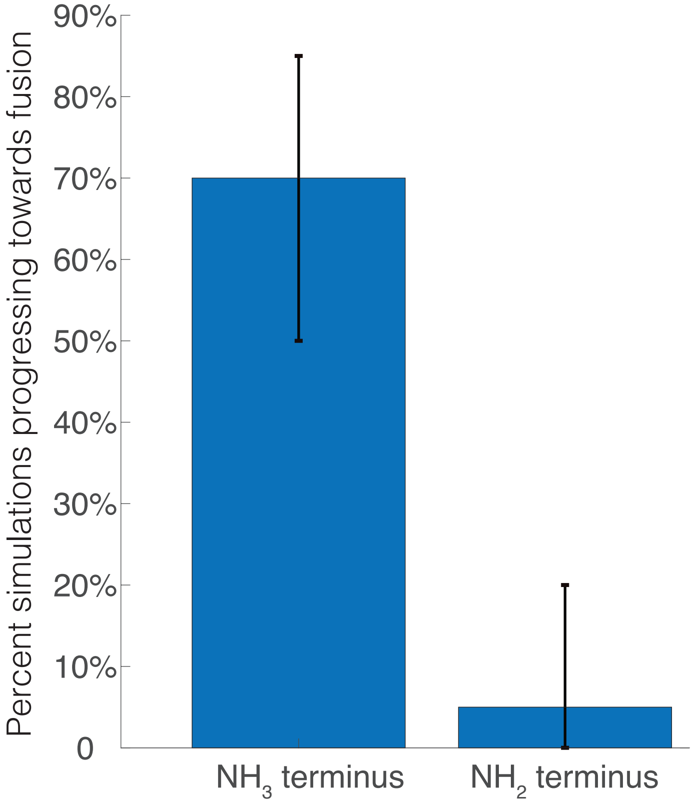


**Figure S6**. **Percentage of simulations with 30-nm proteoliposome and 1930 nm$^2$ bilayer progressing towards fusion**. Simulations were started from a stalk state, and progression was assessed as formation of a terminal hemifusion diaphragm (thinned hemifusion diaphragrm between distal leaflets) or fusion pore formation. Error bars represent 90% bootstrap confidence intervals.

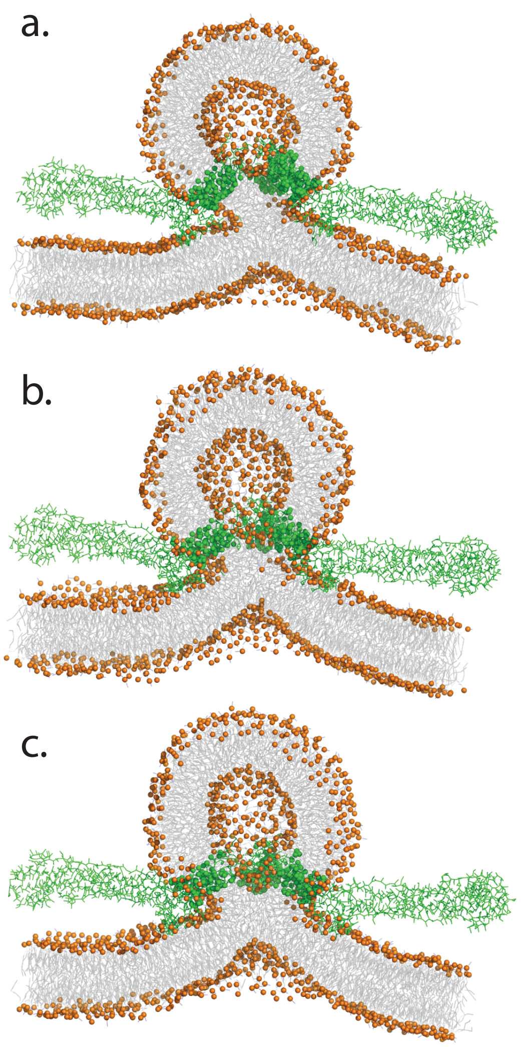

**Figure S7**. **Transmembrane domain pins vesicle inner leaflet into junctional complex**. Renderings of a slice through the proteoliposome inner leaflet are shown for a simulated trajectory where inner leaflet involvement in the junctional complex triggers stalk expansion and hemifusion diaphragm formation. The order of inner leaflet versus target membrane lower-leaflet involvement varies between trajectories. In panel (a), the hemagglutinin transmembrane domains are seen to restrict the inner leaflet to near the junctional complex. This inner leaflet moves into the complex in panel (b), rendered 25 ns later, and the target membrane lower leaflet begins to move upwards. Contact between acyl tails of these two leaflets begins approximately 15 ns later, shown in panel (c).

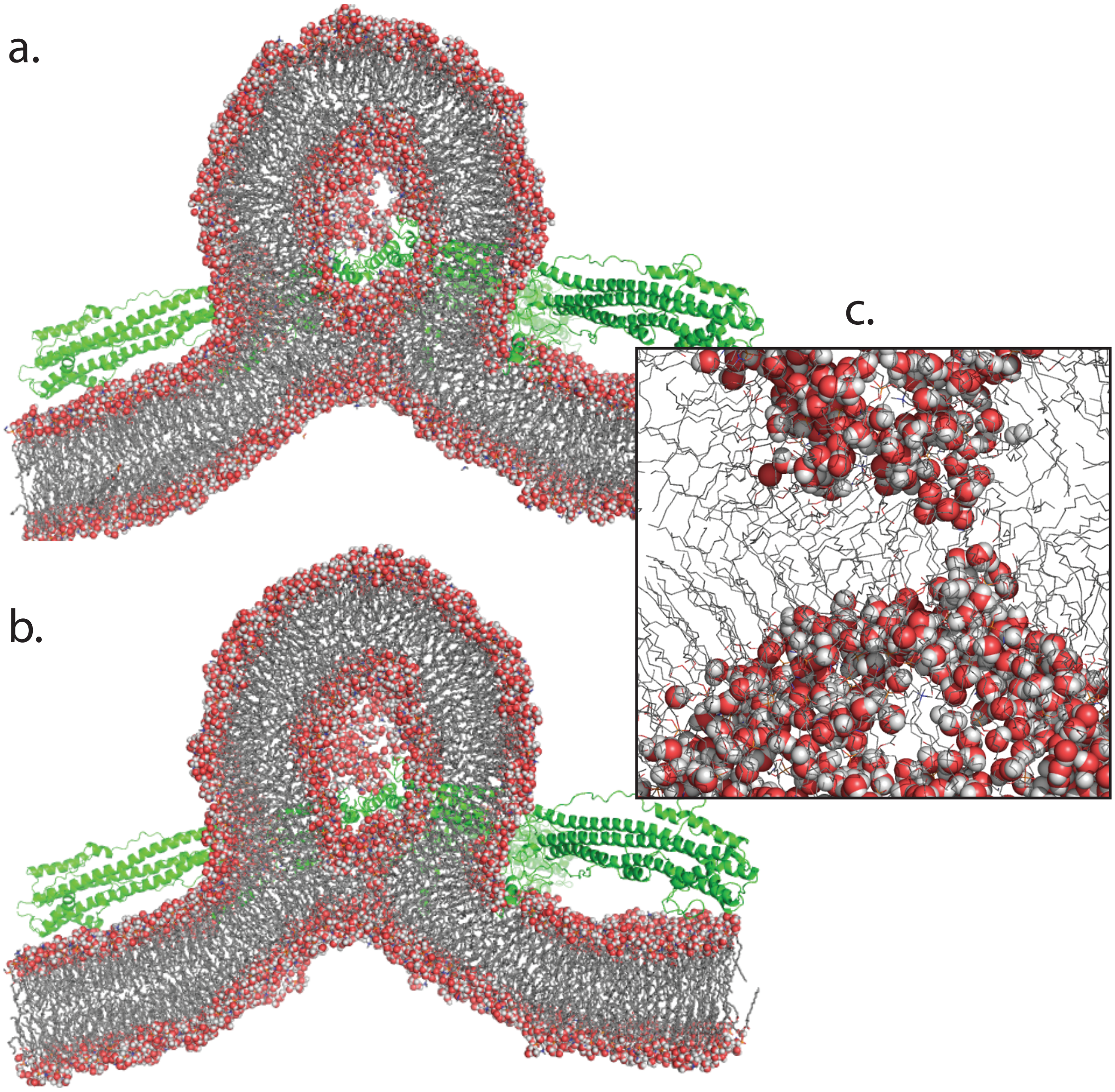


**Figure S8. Renderings of pore opening from atomistic simulations**. Renderings are shown from snapshots of atomistic simulations just before (a) a fusion pore is created and just after (b). Here a fusion pore is defined as continuous water density from the vesicle lumen to the underside of the target bilayer. Water molecules approach each other from both sides of the thinned hemifusion diaphragm (inset, c).

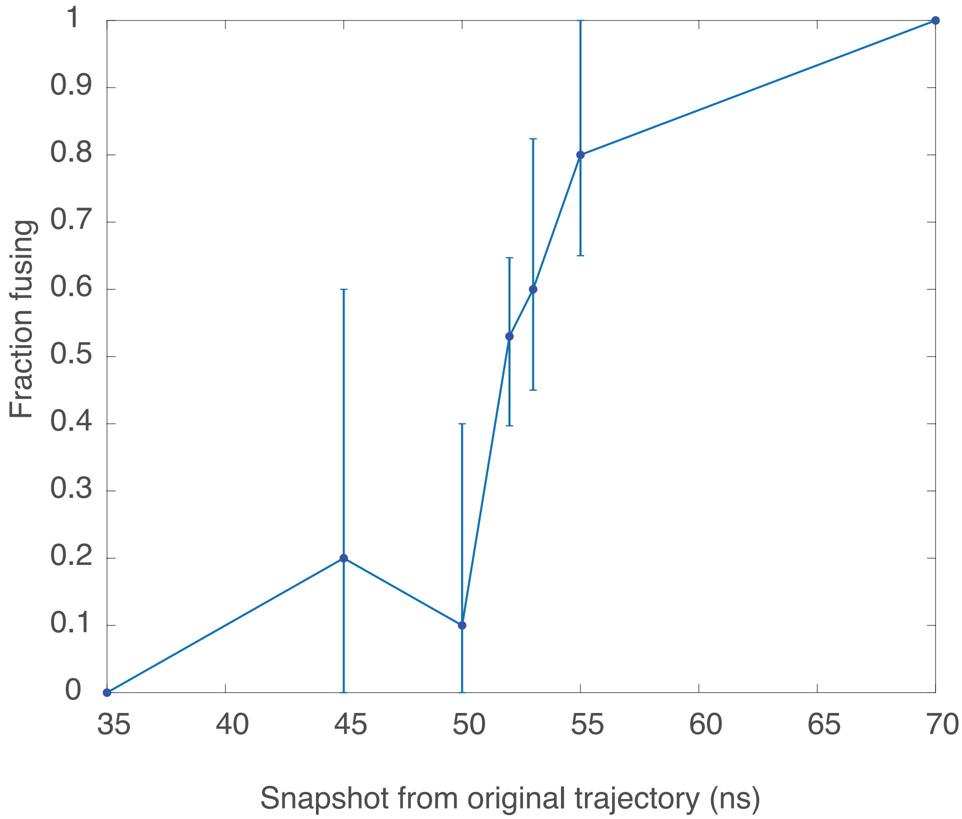


**Figure S9. Gly1 protonation state affects early commitment to stalk expansion and fusion pore opening**. The timing where N-terminal protonation of the fusion peptide exerts its effect was probed by cross-committor analysis (8) as follows. Snapshots were taken from a simulation trajectory with $NH_3$-Gly fusion peptides that ultimately resulted in fusion, and multiple new simulations were started from each snapshot in a traditional committor-analysis procedure. However, these new simulation trajectories were computed using the Hamiltonian for $NH_2$-Gly peptides rather than the original simulation Hamiltonian (hence the term cross-committor). Results of this analysis are plotted, showing that the N-terminal protonation results in an early state change of the system that commits to fusion even if the protonation is removed. Error bars represent 95% bootstrapped confidence intervals.

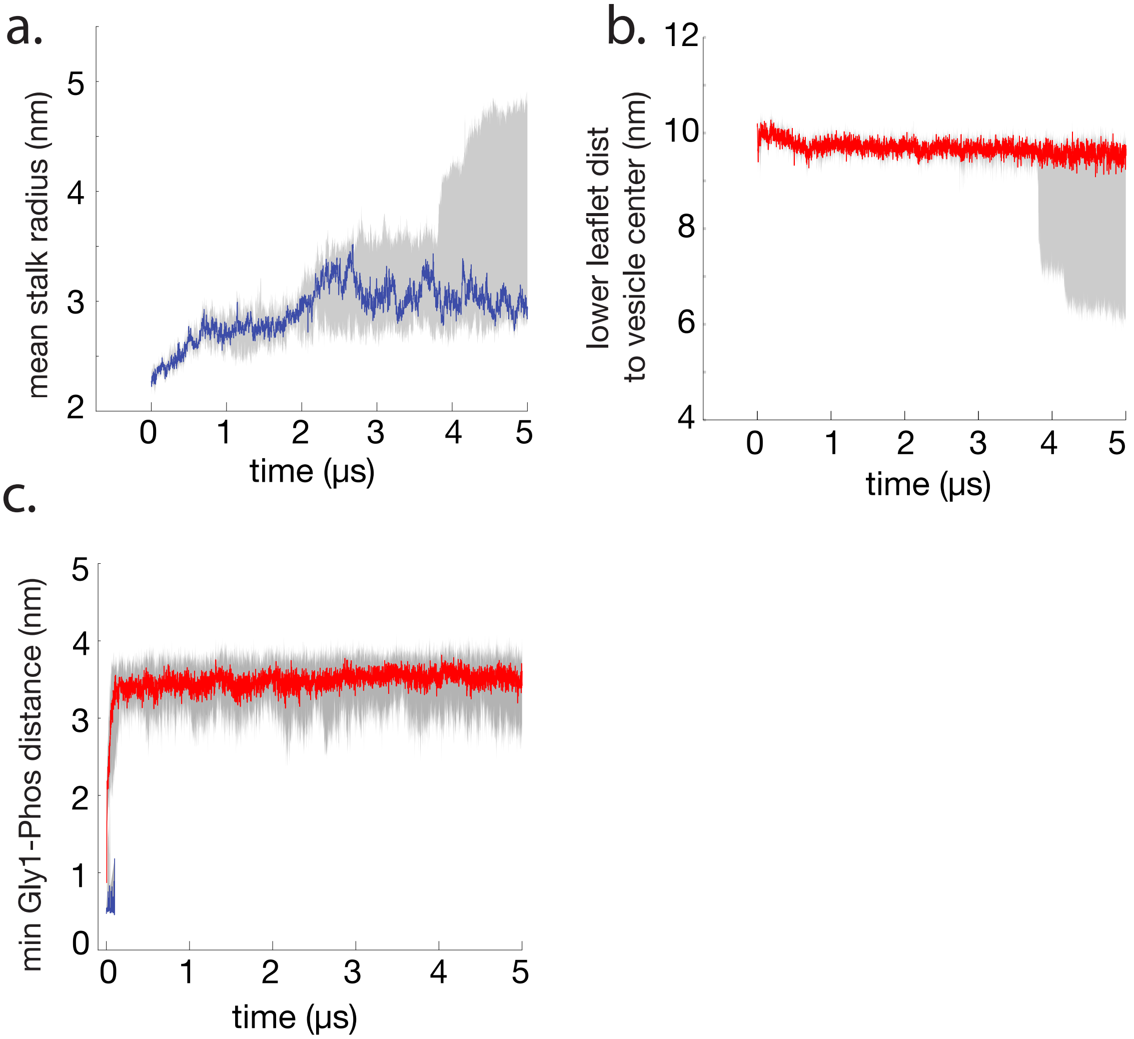


**Figure S10. Lack of stalk expansion and early loss of Gly1-Phosphate contact in $NH_2$-Gly simulations**. Simulations from a stalk state corresponding to Figure 4 were started with deprotonated N-termini. Fusion peptides in these simulations rapidly and irreversibly lost contact with the distal leaflet phosphates, the distal leaflets did not become involved in the junctional complex. Stalk expansion occurred in 4 simulation trajectories, but none of them yielded fusion pore formation on the 5-14 µs timescale of the simulations. In these simulations, stalk expansion occurred much more slowly and was accompanied by lateral pivoting of the hemagglutinin transmembrane domains, consistent with the transmembrane domains "pinning" the proteoliposome inner leaflet into the junctional complex. Simulations with the same fusion peptides and a larger bilayer patch did, however, show fusion (Fig. 5, Fig. S10).

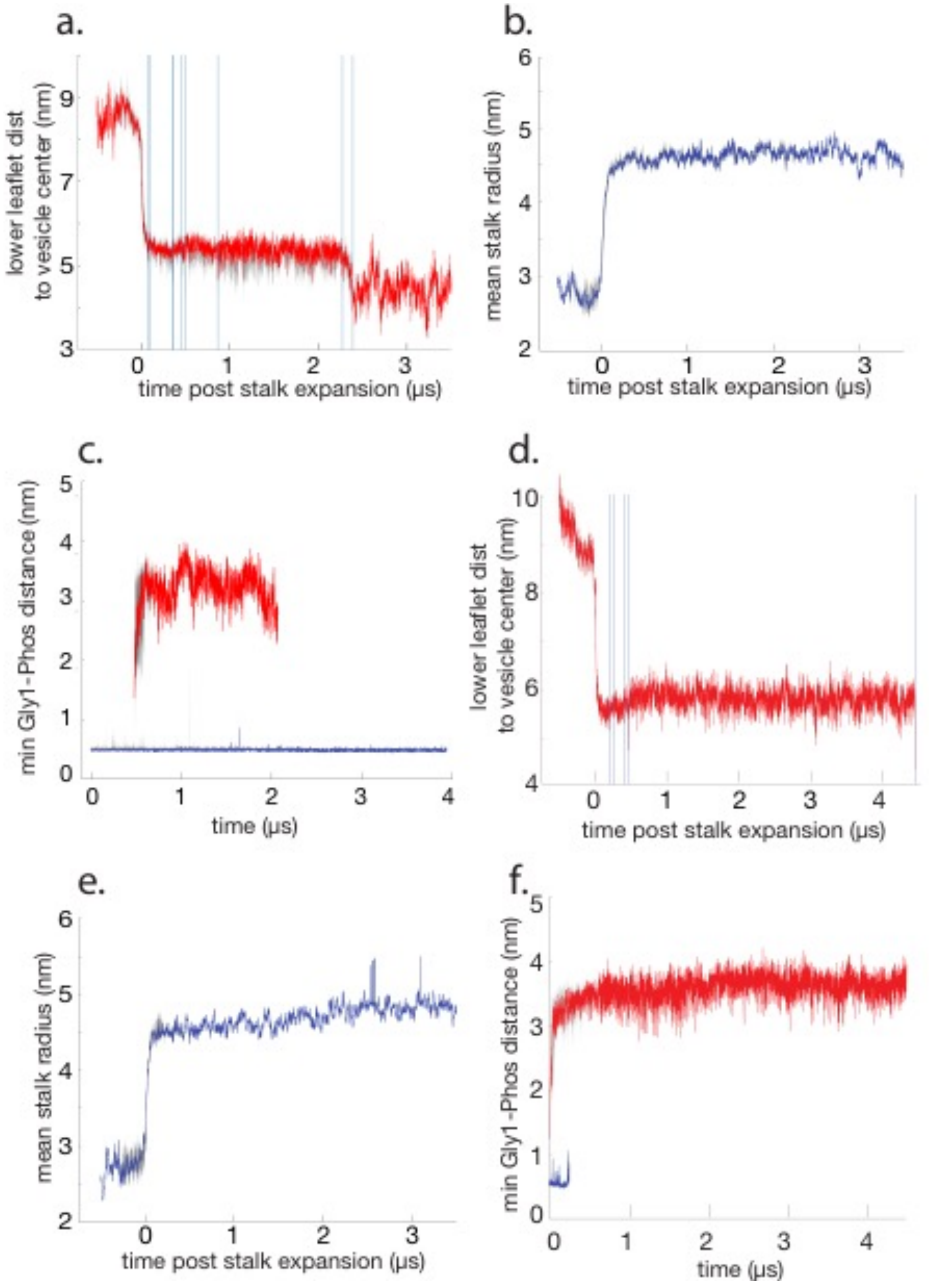


**Figure S11**. **Stalk expansion, distal leaflet approach, and Gly1-Phosphate contact in simulations of larger bilayer patches**. Data from Fig. 5 panels c-e are plotted separately for $NH_3$-Gly simulations (panels a-c, showing stalk size, distal leaflet approach to the junctional complex, and Gly1-lower leaflet phosphate distance) and $NH_2$-Gly simulations (panels d-f) respectively. Labeling schemes are identical to Figure 5; gray represents inter-quartile variation and largely fall within the fluctuation of the median values plotted in blue or red.

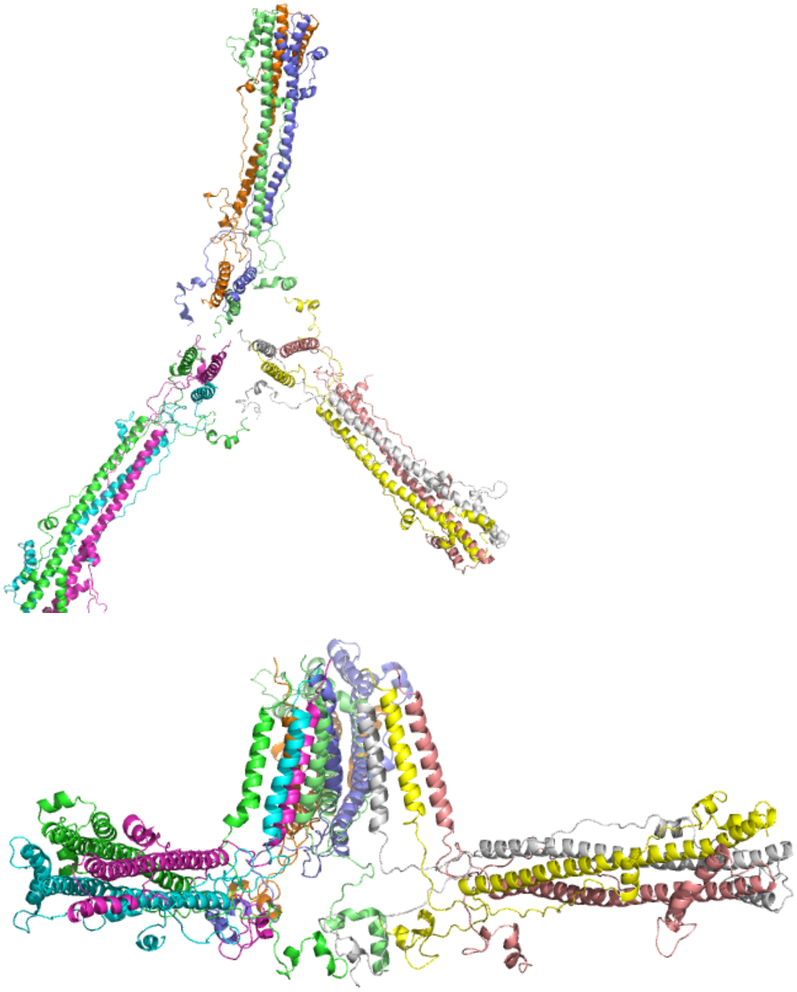

**Figure S12**. **Starting conformations for atomic-resolution simulations**. The protein assembly after initial relaxation is rendered here in top and side views. Individual trimer conformations were generated using Modeller with three structural templates: a previous NMR structure of the fusion peptide residues 1-20, a crystal structure of the soluble ectodomain, and an ideal alpha-helix structure corresponding to the transmembrane domain. Fusion peptides were initially generated at 120º angles with the N-termini facing outwards. In homology modeling, the fusion peptide and transmembrane helix were kept fixed, and the remainder of the trimer structure was generated via homology modeling. These trimers were then inserted into the target bilayer using EPR data for docking and into the vesicle, such that the ectodomains were also facing outward. Both fusion peptide orientation and conformation relaxed over the course of the atomic-resolution runs (see Figs S1, S2 for conformational analyses).

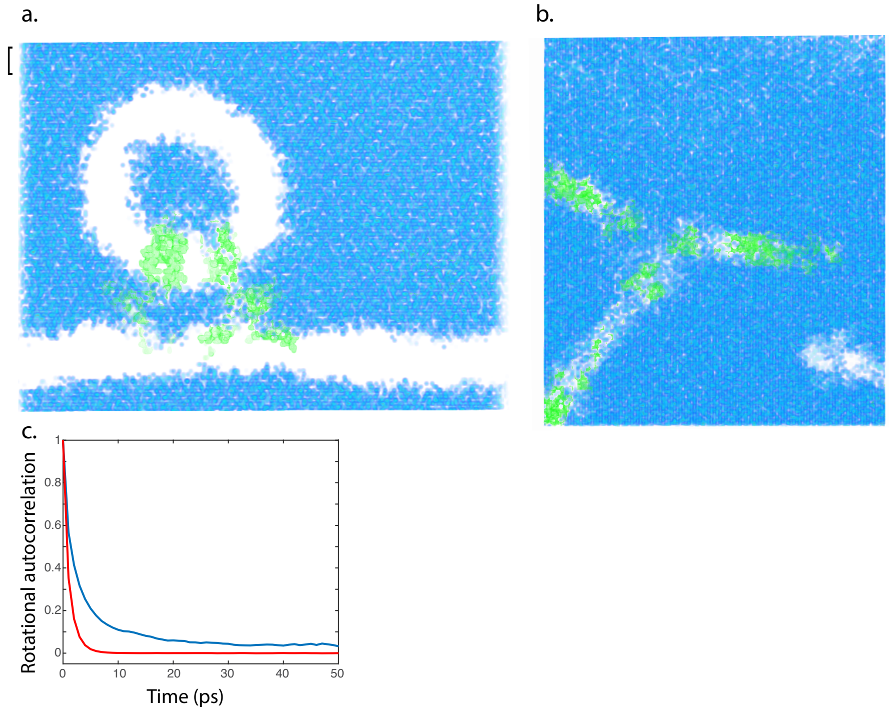


**Figure S13. Water density of starting state for atomic-resolution simulations**. Water density was calculated in voxels 2 Å on a side and rendered in blue for a 20-Å slice through the proteoliposome, shown in panel (**a**). A transverse slice between the proteoliposome and bilayer is shown in panel (**b**). Protein density is rendered in green. The region indicated with a bracket was chosen to estimate bulk TIP3P water density; the average across 18 Å in z (and the full x-y box) in this region was 0.988 g/cm$^3$. Rotational autocorrelation functions are plotted in panel (**c**) for water molecules within 1.5 nm of both the proteoliposome and the bilayer (blue) and water molecules greater than 5 nm from any membrane surface (red) as a bulk reference. Functions plotted are the time autocorrelation of the outer product of the two OH bond vectors. Self-diffusion coefficients for the interfacial water molecules were estimated at 1.9 ± 0.24 x $10^{-5}$ cm$^2$/s, ~3 fold slower than bulk TIP3P water at 310 K.

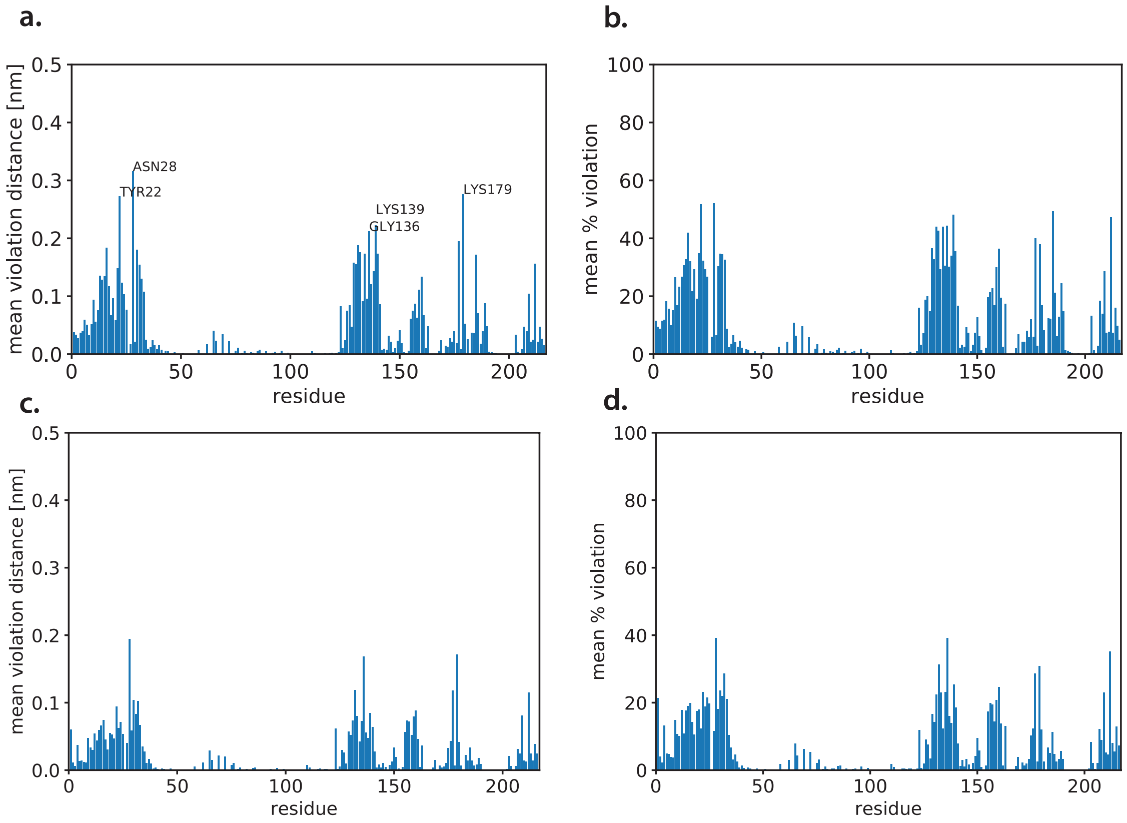


**Figure S14. Preservation of coarse-grained conformational restraints during atomic-resolution simulations**. Coarse-grained simulations in MARTINI use elastic restraints to preserve secondary structure; these were automatically generated using the standard MARTINI protocol (9). Deviations from the restraint equilibrium values were calculated in atomistic simulations and are plotted here for simulations with protonated N-termini (a, b) and neutral N-termini (c, d) respectively. Percent deviation was calculated using a 0.23 nm cutoff, the radius of a coarse-grained bead. Some deviation is expected, and the only substantial deviations measured were in flexible linker regions.

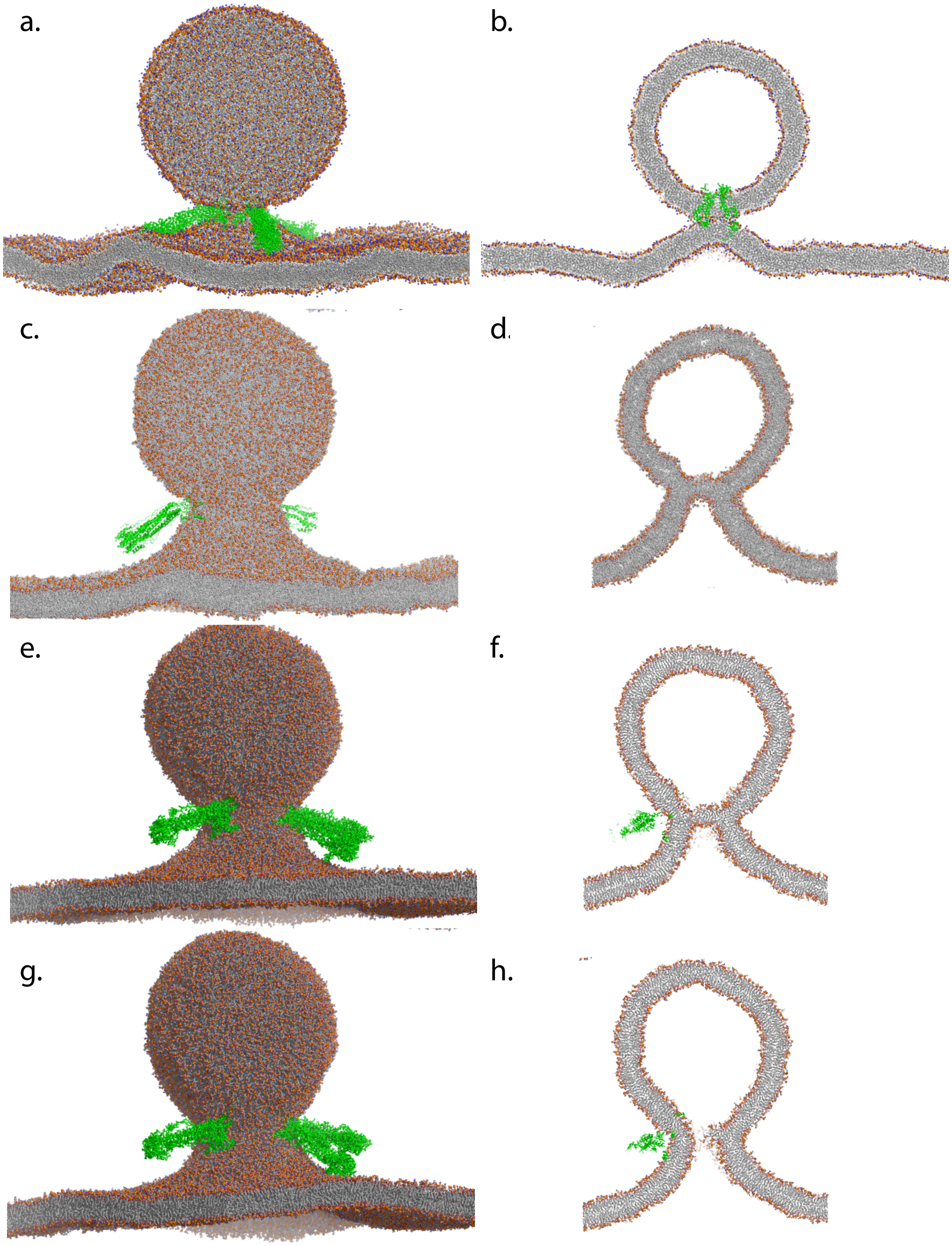


**Figure S15. Renderings of fusion snapshots between 30-nm proteoliposome and target bilayer**. Shown are snapshots of the full system minus solvent at time of stalk formation and initiation of coarse-grained simulation (a), conversion to atomic resolution shortly before pore formation (c), time of pore formation (e), and 6 ns following pore formation (g). 20-nm slices through the proteoliposome are also rendered at the corresponding time-points (b, d, f, h).

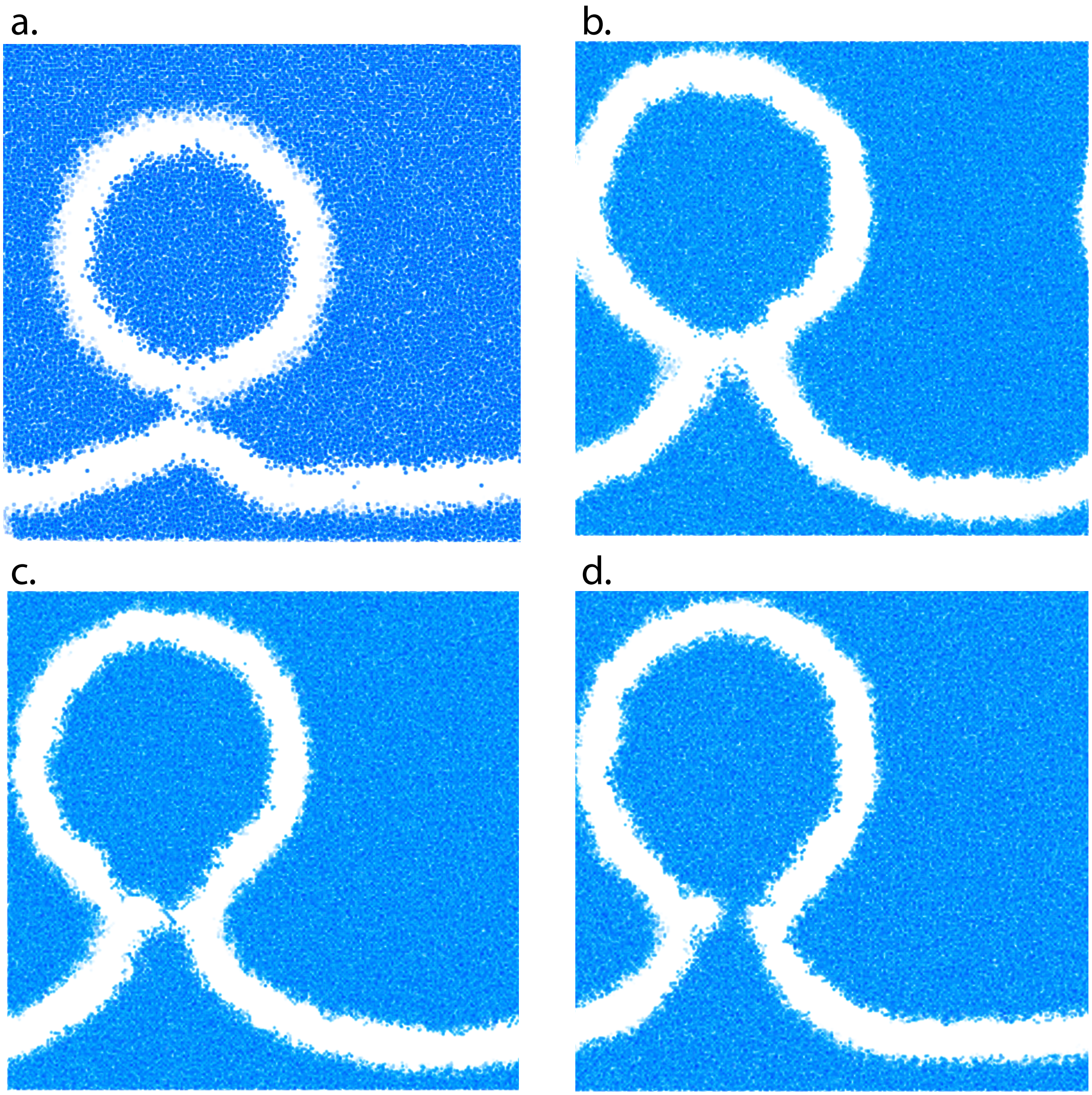


**Figure S16. Water density in simulations of 30-nm proteoliposome fusion**. Shown are renderings of water density at time of stalk formation and initiation of coarse-grained simulation (a), conversion to atomic resolution shortly before pore formation (b), time of pore formation (c), and 6 ns following pore formation (d).

a. **Atomistic simulations from a docked state**

| **Start state** | **Parameters** | **# runs** | **Run lengths** |
|---|---|---|---|
| Docked proteoliposome | AMBER99SB-ILDN, $NH_2$-Gly | 50 | Min 203 ns, max 500 ns, median 500 ns |
| Docked proteoliposome | AMBER99SB-ILDN, $NH_3$-Gly | 18 | Min 122 ns, max 378 ns, median 238 ns |

b. **Coarse-grained simulations from an atomistic stalk state**

| **Start state** | **Parameters** | **# runs** | **Run lengths** |
|---|---|---|---|
| Initial Stalk | MARTINI v2.2, $NH_3$-Gly, POPE:POPC | 10 | Min 273 ns, max 16 μs, median 2.2 μs |
| Initial Stalk | MARTINI v2.2, $NH_2$-Gly, POPE:POPC | 10 | Min 5.4 μs, max 14.7 μs, median 8.2 μs |
| Initial Stalk | MARTINI v2.2, $NH_3$-Gly, POPC | 10 | Min 2.8 μs, max 26.5 μs, median 21.1 μs |
| Initial Stalk | MARTINI v2.3, $NH_3$-Gly, 25 DOPE:75 POPC | 10 | Min 675 ns, max 4.7 μs, median 2.2 μs |
| Initial Stalk | MARTINI v2.3, $NH_2$-Gly, 25 DOPE:75 POPC | 10 | Min 765 ns, max 6.0 μs, median 5.4 μs |

c. **Coarse-grained simulations with a larger target bilayer**

| **Start state** | **Parameters** | **# runs** | **Run lengths** |
|---|---|---|---|
| Initial Stalk, larger bilayer | MARTINI v2.2, $NH_3$-Gly, POPE:POPC | 15 | Min 228 ns, max 10.6 μs, median 1.1 μs |
| Initial Stalk, larger bilayer | MARTINI v2.2, $NH_2$-Gly, POPE:POPC | 15 | Min 463 ns, max 7.5 μs, median 3.6 μs |

d. **Cross-committor simulations from a coarse-grained $NH_3$-Gly trajectory**

| **Start state** | **Parameters** | **# runs** | **Run lengths** |
|---|---|---|---|
| 35 ns after stalk formation with CG $NH_3$-Gly | MARTINI v2.2, $NH_2$-Gly, POPE:POPC | 5 | Min 1.5 μs, max 2.2 μs, median 1.8 μs |

| | | | |
|---|---|---|---|
| 45 ns after stalk formation with CG $NH_3$-Gly | MARTINI v2.2, $NH_2$-Gly, POPE:POPC | 5 | Min 2.3 µs, max 2.6 µs, median 2.4 µs |
| 50 ns after stalk formation with CG $NH_3$-Gly | MARTINI v2.2, $NH_2$-Gly, POPE:POPC | 10 | Min 2.0 µs, max 2.2 µs, median 2.1 µs |
| 52 ns after stalk formation with CG $NH_3$-Gly | MARTINI v2.2, $NH_2$-Gly, POPE:POPC | 60 | Min 1.0 µs, max 12.6 µs, median 2.6 µs |
| 53 ns after stalk formation with CG $NH_3$-Gly | MARTINI v2.2, $NH_2$-Gly, POPE:POPC | 20 | Min 1.2 µs, max 7.6 µs, median 2.7 µs |
| 55 ns after stalk formation with CG $NH_3$-Gly | MARTINI v2.2, $NH_2$-Gly, POPE:POPC | 5 | Min 4.0 µs, max 5.0 µs, median 4.5 µs |
| 70 ns after stalk formation with CG $NH_3$-Gly | MARTINI v2.2, $NH_2$-Gly, POPE:POPC | 5 | Min 1.7 µs, max 4.3 µs, median 2.2 µs |

**e. Atomistic simulations of pore formation**

| **Start state** | **Parameters** | **# runs** | **Run lengths** |
|---|---|---|---|
| 10ns prior to pore opening in CG $NH_3$-Gly simulations | CHARMM36, $NH_3$-Gly | 19 | Min 5.2 ns, max 96.1 ns, median 86.5 ns |
| 10ns prior to pore opening in CG $NH_3$-Gly simulations | CHARMM36, $NH_2$-Gly | 11 | Min 45.4, max 71.6, median 68.4 ns |
| Late hemifusion diapgragm in CG $NH_3$-Gly simulations with larger vesicle and bilayer | CHARMM36, $NH_3$-Gly | 10 | Min 27.1 ns, max 52.1 ns, median 39.1 ns |

**Table S1. Simulations and run lengths used in analyses of hemagglutinin-mediated fusion.** Each set of start states, simulation parameters, runs, and lengths is tabulated here. Minimum, maximum, and median values are reported for lengths.

**Supplementary Methods**.

*Atomic-resolution simulations*. Simulations from the docked state were performed using a starting conformation described in the Methods and available as part of a Dryad package to accompany this paper. Simulations were run in Gromacs (10) using the AMBER99SB-ILDN force field (11, 12) and lipid parameters based on the Berger parameter set (13) and previously published (14). Simulations were solvated in TIP3P water with 150 mM NaCl using standard Gromacs utilities. Simulations used a 1.2 nm cutoff for direct-space interactions and treated long-range electrostatics with Particle Mesh Ewald (15). Temperature was maintained at 310º K using a velocity-rescaling thermostat (16) with a 5-ps relaxation constant, and pressure was maintained at 1 bar using a Berendsen barostat applied semi-isotropically with a 10-ps relaxation constant. For AMBER simulations, bonds were constrained using LINCS, and a 4-fs timestep was used. For CHARMM simulations, a 2-fs timestep was used. Full parameter files are available as part of the Dryad package. Atomic-resolution simulations of pore opening and one set of simulations from the docked state used CHARMM36 protein and lipid parameters (17, 18) made available by MacKerell and co-workers instead (again run using the Gromacs engine).

*Coarse-grained simulations*. Coarse-grained simulations used MARTINI v2.2. Production parameters were as follows, largely following standard practice for MARTINI. Direct-space interactions were shifted to zero at 1.2 nm, and long-range electrostatics were approximated using a reaction-field treatment with a dielectric constant of 15. A 20-fs timestep was used, and temperature was maintained at 310º K using a velocity-rescaling thermostat with a 12-ps relaxation constant. Except as noted, coarse-grained simulations were run in the NVT ensemble to avoid the possibility of inducing fusion via exogenous changes to membrane tension. Use of the NVT ensemble may impede fusion relative to a tension-free bilayer, since fusion involves substantial membrane bending, but the fusion events observed in these ensembles are known to be free of barostat-induced artifacts. Control simulations performed using the NPT ensemble showed fusion similar to the NVT data reported here. We have elected not to include time-correction factors between coarse-grained and atomistic simulations in our nomenclature; elapsed time in MARTINI coarse-grained simulations has been estimated to correspond to 3-4x that amount in atomistic simulations.

*Generation of larger bilayer patches*. Initial coarse-grained starting conformations in the stalk state were tiled in a 3x3 fashion and then a larger hexagonal prism was cut out surrounding the central stalk structure, essentially replicating the "bulk" lipid patch away from the stalk region. This generated a stalk structure with total bilayer 33.7 nm on a side in the periodic cell from the original one with bilayer 25.8 nm on a side. During energy minimization, positional restraints were applied to all lipids, and the new bilayer edges were pulled together (since they did not precisely align due to membrane undulation) using a harmonic potential with force constant 300 kJ $mol^{-1}$ $nm^{-2}$. The resulting structure was then equilibrated for 250 ps prior to production simulation with either a protonated or an unprotonated N-terminal glycine as designated. Starting conformations are available as part of the Dryad package to accompany this paper.

*Generation of larger vesicle:bilayer stalk structures*. The 33.7-nm bilayer generated above was expanded a second time in the same fashion, yielding a bilayer with lateral dimensions 47.2 nm on a side. Equilibration was performed as described above. A larger vesicle was generated using the MARTINI Maker package (19), specifying a vesicle radius (here defined as center-to-bilayer-midplane distance) of 13 nm, approximately twice the radius of the initial vesicle, and a composition of 75:25 POPE:POPC. This vesicle was equilibrated in Gromacs (10) using the protocol recommended by the MARTINI Maker package authors. The equilibrated vesicle was then fit to the stalk structure that served as the starting state for the original coarse-grained simulations of small vesicle-bilayer fusion as follows. Vesicles were aligned using rigid-body fitting to the vesicle-bilayer contact patch in the stalk structure, selected as phosphate groups in a z-slice of 1.3 nm corresponding to this contact patch and a corresponding 1.3-nm z-slice in the larger vesicle. Within this slice, lipids from the original stalk structure were maintained, and the remainder of the small vesicle was replaced with the large vesicle. This protocol was designed in analogy to that previously performed for nascent fusion pores (20). This hybrid stalk structure was then equilibrated using position restraints on 1) the phosphate groups of all lipids involved in the stalk, 2) the hemagglutinin fusion peptides, and 3) the hemagglutinin transmembrane domains. Position restraints were applied for NVT equilibration runs totaling 600ps and restraint force constants varied between 1000 and 0 kJ $mol^{-1}$ $nm^{-2}$. After that, production runs were initiated using the same simulation parameters as for coarse-grained simulations of the initial vesicle-bilayer complex. Simulations were performed using either protonated or unprotonated fusion peptide N-termini, and control simulations were performed using the 47.2-nm bilayer but the original vesicle. Three of these production runs were converted to atomic resolution shortly before fusion pore opening using the procedure described in the Methods, and three atomic-resolution simulations were performed per starting state for a total of 9 atomic-resolution simulations of fusion pore formation between a 13-nm proteoliposome and a 47.2-nm bilayer.

*Analysis of stalk formation and pore opening*. Stalk formation and fusion pore opening were measured using alpha-complexes, a topological construct used previously to measure fusion (21). Essentially, alpha complexes use balls of radius alpha to connect point clouds into triangulated surfaces. We used the number of connected components of proximal-leaflet (bilayer upper leaflet and vesicle outer leaflet) acyl tail groups to monitor stalk formation and the number of connected components of distal-leaflet (bilayer lower leaflet and vesicle inner leaflet) head groups to monitor pore formation. Alpha values used were 0.3 nm for stalk formation and 0.5 nm for pore formation; analysis code is available from https://github.com/kassonlab/fusion-shape-analysis in conjunction with publication of this paper.
